# Confounding factors in targeted degradation of short-lived proteins

**DOI:** 10.1101/2024.02.19.581012

**Authors:** Vesna Vetma, Laura Casarez-Perez, Ján Eliaš, Andrea Stingu, Anju Kombara, Teresa Gmaschitz, Nina Braun, Tuncay Ciftci, Georg Dahmann, Emelyne Diers, Thomas Gerstberger, Peter Greb, Giorgia Kidd, Christiane Kofink, Ilaria Puoti, Valentina Spiteri, Nicole Trainor, Yvonne Westermaier, Claire Whitworth, Alessio Ciulli, William Farnaby, Kirsten McAulay, Aileen B. Frost, Nicola Chessum, Manfred Koegl

## Abstract

Targeted protein degradation has recently emerged as a novel option in drug discovery. Natural protein half-life is expected to affect the efficacy of degrading agents, but to what extent it influences target protein degradation has not been systematically explored. Using mathematical modelling of protein degradation, we demonstrate that the natural half-life of a target protein has a dramatic effect on the level of protein degradation induced by a degrader agent which can pose significant hurdles to screening efforts. Moreover, we show that upon screening for degraders of short-lived proteins, agents that stall protein synthesis, such as GSPT1 degraders and generally cytotoxic compounds, deceptively appear as protein degrading agents. This is exemplified by the disappearance of short-lived proteins such as MCL1 and MDM2 upon GSPT1 degradation and upon treatment with cytotoxic agents such as doxorubicin. These findings have implications for target selection as well as for the type of control experiments required to conclude that a novel agent works as a bona fide targeted protein degrader.

## Introduction

In the last decade, targeted protein degradation has been established as a new therapeutic modality in small molecule drug discovery ^1–6^. Targeted degradation can be achieved by inducing the proximity of a ubiquitinating enzyme to a protein of interest (POI). This results in the covalent modification of the POI with multi-ubiquitin chains, which are recognized by the proteasome, the cell’s major protease, and results in the protein degradation. Proximity-inducing small molecules have been classified as molecular glues or PROTACs (proteolysis targeting chimeras). Whilst glues work by reshaping the surface of the ligase to enhance binding to the POI, PROTACs consist of separate moieties binding to the ligase and the POI, connected by a linker ^7^. Molecular glues have been discovered mostly by serendipity with respect to the target, the ligase or both ^8–14^. In contrast, the bipartite structure of PROTACs facilitates, in principle, their rational design for any POI for which small molecule binders exist.

In drug discovery, PROTACs may be an option if binders to the target do not modulate its activity, or if modulation of the activity has no or insufficient pharmaceutical consequences. Genetic studies have shown that this is the case for CRAF (RAF1). CRAF is a component of the Ras-MAPK signal transduction pathway. This pathway is frequently altered in cancer by mutations decoupling pathway activation from external stimuli, e.g., by mutations activating Ras-dependent signaling ^15^. In genetic mouse models of KRAS-driven lung cancers, loss of CRAF causes tumor regression ^16,17^, whilst loss of CRAF kinase activity has no effect ^18^. Therapeutic exploitation would therefore require removal of the whole CRAF protein, i.e., require CRAF degradation. However, no CRAF degrading small molecules have been reported to date.

An important point to consider for PROTAC projects is the stability of the target. The natural half-life of a PROTAC target has a significant effect on its ability to be degraded using a targeted approach ^19,20^. We describe herein a mathematical model predicting the maximal degradation that can be achieved with a PROTAC depending on the natural half-life of the target. Drawing from experience in a project targeting CRAF we show that degraders of GSPT1, a known off-target for CRBN-based PROTACs, will artificially appear as degraders of short-lived proteins due to their effect on protein synthesis. In addition, cytotoxic effects can cause a drop in the level of short-lived proteins, resulting in the false classification of such agents as degraders. Many of the control experiments conventionally used in the field, such as proteasome inhibition, fail to differentiate such artifacts from bona fide degraders. The avoidance of these pitfalls in drug discovery projects is discussed.

## Results

### Calculating the effect of protein half-life on targeted protein degradation

Intuitive consideration would suggest that it should be easier to degrade a long-lived protein than a short-lived one using targeted degraders. Nevertheless, we wanted to capture this assumption in precise terms. A mathematical model was therefore established that incorporates the initial speed of degradation. Let *P*(*t*) be the amount of the protein of interest at time *t*, then the rate of change

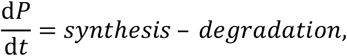

is a conservation equation for the protein amount. The simplest model has constant synthesis and degradation proportional to *P*, that is,

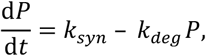

which implies

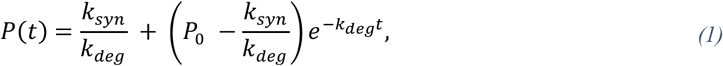

where *k*_*syn*_ and *k*_*deg*_ are positive constants and *P*_0_ = *P*(*t* = 0) is the initial amount of the protein. Independently of *P*_0_ the protein will stabilize at the value *P*_*SS*_ = *k*_*syn*_ /*k*_*deg*_. Moreover, *k*_*deg*_ = *ln*(*2*)/*t*_1/2_ where *t*_1/2_ is the half-life of the protein.

Targeted protein degradation by PROTACs can be modelled by adding a drug dependent degradation term into Equation (1),

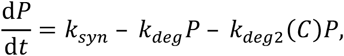

with a reasonable assumption for the initial protein amount *P*_0_ = *P*_*SS*_. The drug concentration *C* is either constant (e.g., in *in vitro* experiments) or a time dependent curve following a pharmacokinetic profile. If *C* is constant, then the protein fraction left after degradation at any time *t* and for any *C* is given by

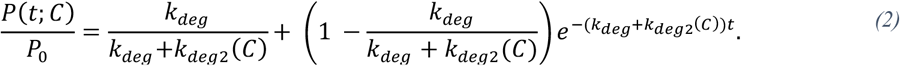

The form of *k*_*deg2*_ (*C*) necessitates modelling the situation with which we are concerned. *In vitro* degradation data not showing any hook effect can be well modelled with a sigmoid function *k*_*deg2*_ (*C*) = *k*_*inact*_ *C*/(*K*_*i*_ + *C*) where *k*_*inact*_ and *K*_*i*_ are positive constants (Bartlett and Gilbert 2022).

### Prediction of degradation of artificially short-lived proteins

Degradation assays based on the use of protein fusions with tags that allow quick and easy quantification of protein abundance, such as the HiBit-tag ^21^, are very helpful for the characterization of protein degraders. We intended to use the HiBit tag in a project looking for CRAF degrading PROTACs. HiBit-fusion proteins were stably expressed using retroviral transduction. However, when we assessed the stability of the HiBit fusion proteins upon inhibition of protein synthesis, we were surprised to see a significantly quicker decay of the HiBit-fusions than of the endogenous protein (Figure 1A). The effect was particularly pronounced for a mutant version of CRAF (CRAF S259A) containing a mutation in the phosphorylation site required for repressive binding to 14-3-3 proteins. The effect of the HiBit-tag was also apparent for BRAF, particularly for BRAF V600E, but not for N-terminal fusions of BRAF or CRAF with other short tags, such as the FLAG or V5 tag, or with full-length luciferase (Figure 1B). Increased degradation rates were also seen for other HiBit-tagged proteins tested, such as SMARCA2 and SMARCA4 (Figure 1C). Note that other proteins, such as KRAS, were not destabilized by the tag ^22^. Thus, while the HiBit-tag does not generally make proteins short-lived, it points towards the necessity of carefully assessing the half-lives of fusion proteins in comparison with the endogenous proteins.

**Figure 1.**
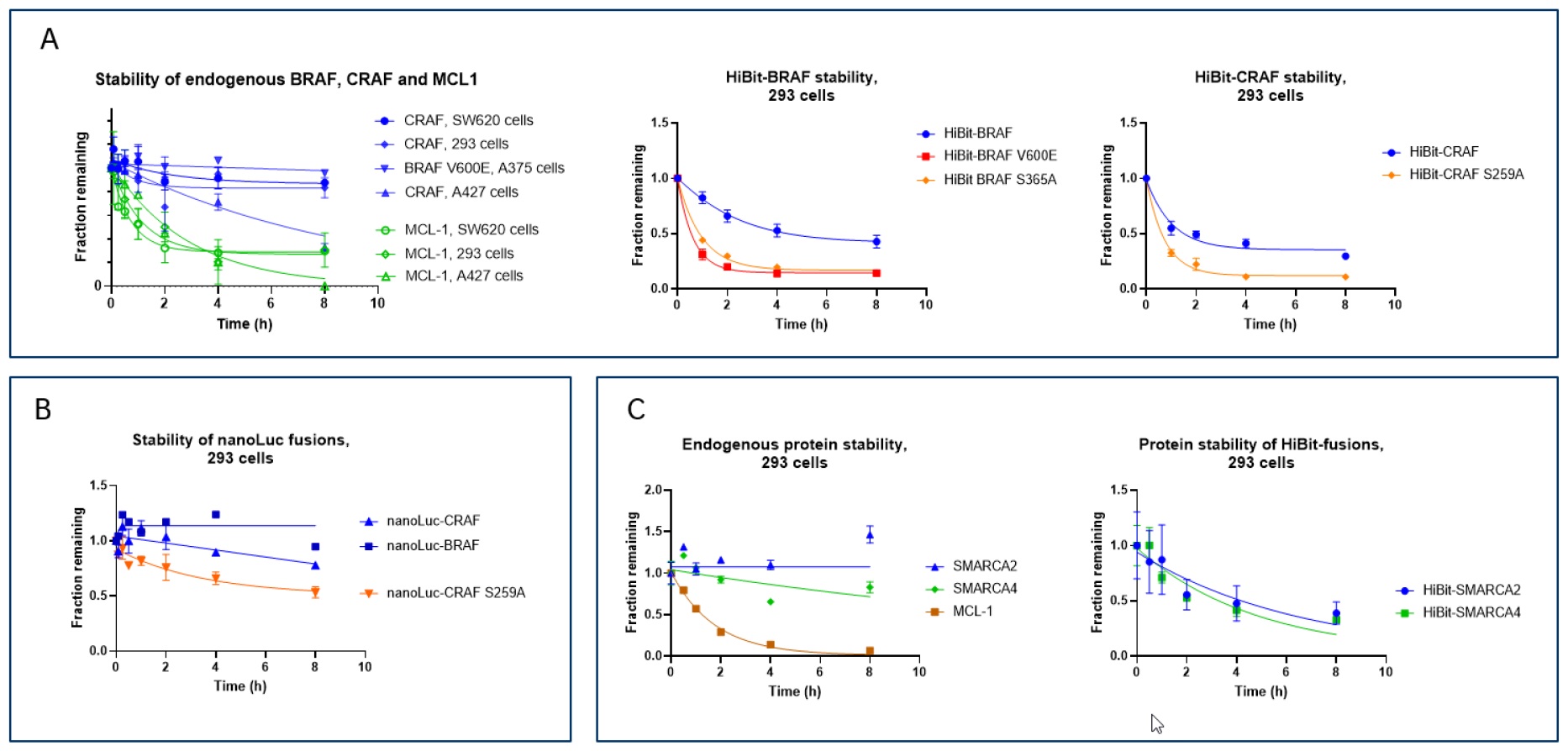
HiBit tagging can render proteins short-lived. Decay curves of the indicated proteins after blockage of protein synthesis by cycloheximide were measured in the indicated cell lines. Protein levels were determined by capillary electrophoresis for endogenous proteins and by HiBit or luciferase assays for HiBit and nanoLuc fusions. **A**, Decay curves of endogenous and HiBit-tagged RAF proteins and of endogenous MCL1 as an example for a short-lived protein. **B**, Stability of nanoLuc fusions of BRAF and CRAF. **C**, Decay curves of endogenous and HiBit-tagged SMARCA2 and -4 and of endogenous MCL1.

Our inadvertent discovery that the HiBit-tag can render naturally long-lived proteins short-lived allowed us to test the mathematical model. Equation (2) with *k*_*deg2*_ (*C*) = *k*_*inact*_*C*/(*K*_*i*_ + *C*) was applied to approximate the degradation of endogenous SMARCA2, which has a half-life exceeding 24 hours. We then used the model to ask the question: how would the dose-response curve of the same PROTAC change if SMARCA2 was short-lived? The result is shown in Fig. 2: More short-lived proteins would be degraded to a much lower extent, as intuitively expected. Notably, proteins with a half-life of less than 2 hours would be minimally degraded, with maximal degradation falling into the level of noise of most degradation assays. Based on the degradation of endogenous SMARCA2 the model correctly approximates the less efficient degradation observed for the more short-lived HiBit-SMARCA2, which has a half-life of approximately 4 hours. (Figure 1C). This emphasizes the fact that a half-life of a protein of interest should be known prior to estimating *k*_*inact*_ and *K*_*i*_ and highlights the need for higher *k*_*inact*_ values to achieve significant maximal degradation of short-lived proteins.

**Figure 2.**
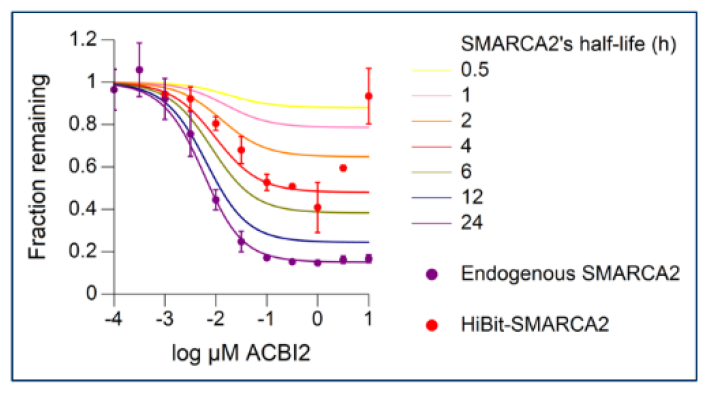
The half-life of a protein affects its degradation by a PROTAC. Equation (2) with *k*_*deg2*_ (*C*) = *k*_*inact*_ *C*/(*K*_*i*_ + *C*) was used to model endogenous SMARCA2 degradation data by ACBI2 PROTAC assuming SMARCA2 to have 24h half-life (purple dots – measured remaining fraction of SMARCA2, purple line – modelled degradation curve). Equation (2) with the fixed kinetic parameters *k*_*inact*_ = 0.19 /h and *K*_*i*_ = 20 nM from the modelling was used to simulate degradation responses for multiple shorter half-lives of SMARCA2 (colored lines other than purple). HiBit-SMARCA2 degradation data (red dots) were then plotted on top of the predicted curves.

### Stalling of protein synthesis via GSPT1 degradation causes a drop in the levels of short-lived proteins

In our attempts to discover PROTACs that degrade CRAF, a library of 6500 compounds was generated by fusing a total of 20 different RAF binders to several E3 ligase binders, including binders of CRBN, VHL, IAP and MDM2. Screening this library using the short-lived HiBit-CRAF S259A construct yielded several hits (Figure 3) that reproducibly reduced the level of HiBit-CRAF S259A. These hits also caused a reduction of endogenous CRAF, albeit to smaller extents, as exemplified for ACBI-8451 (Figure 4A). In time course experiments, the observed degradation turned out to be slow and displayed a conspicuous delay in its onset, suggesting that the observed effects on CRAF may depend on the prior completion of another process (Figure 4B).

**Figure 3.**
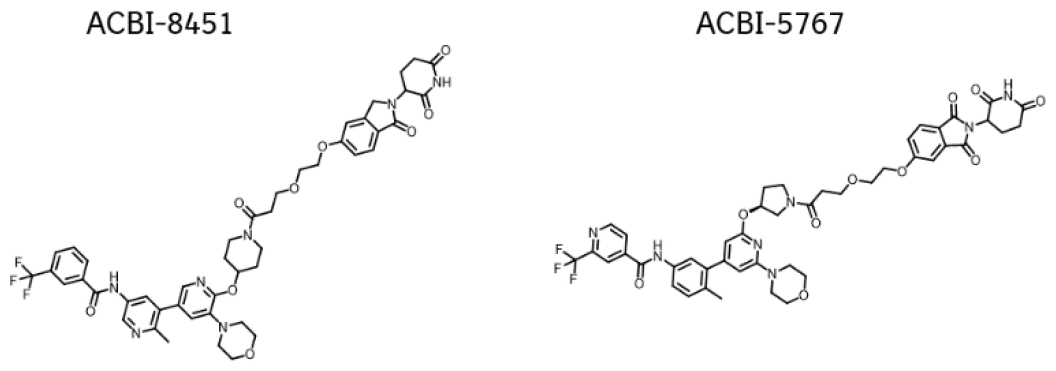
Chemical structures of ACBI-8451 and ACBI-5767.

**Figure 4.**
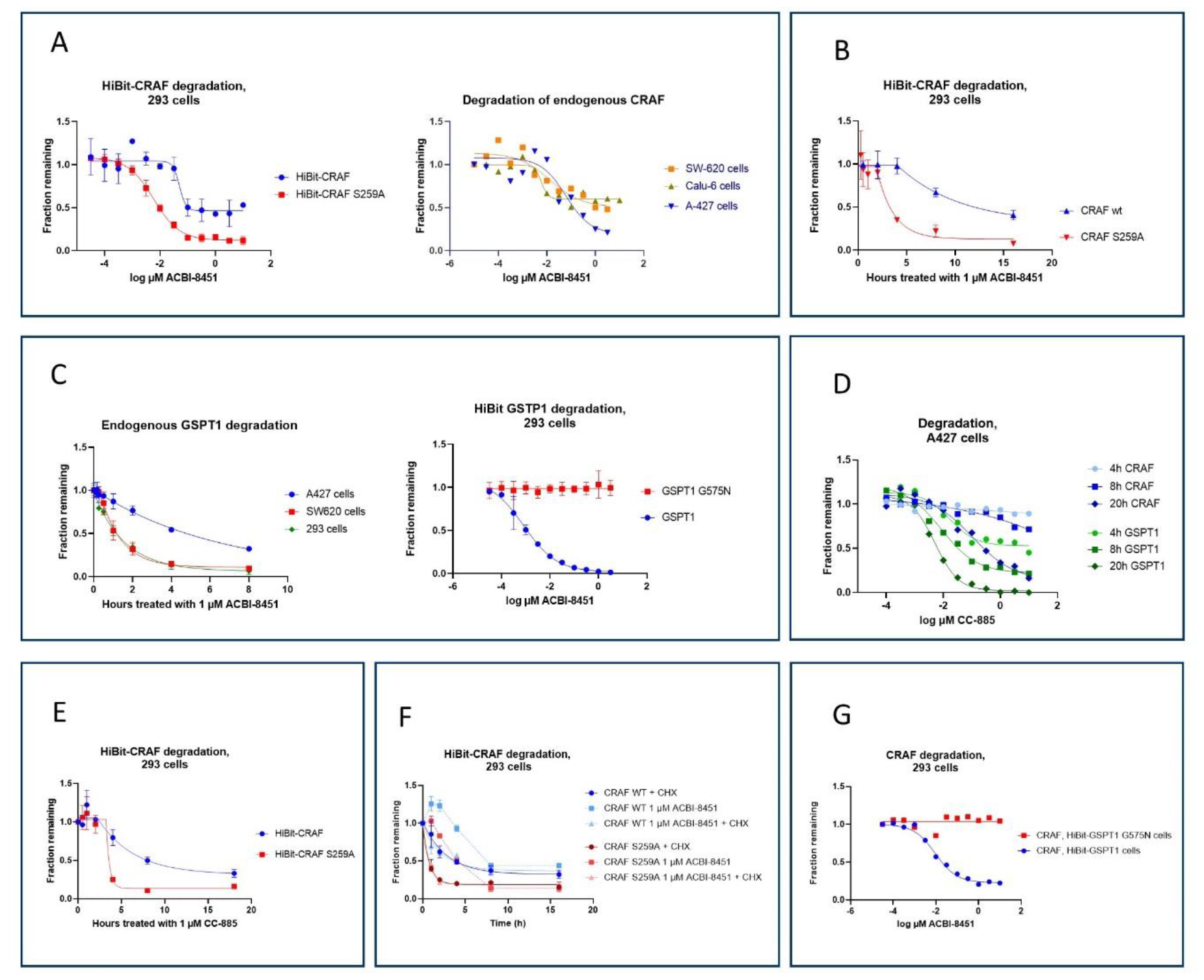
Degradation of CRAF S259A is an artifact brought about by the degradation of GSPT1. **A**, Degradation of CRAF HiBit fusions in 293 cells and of endogenous CRAF in A427, SW620 and Calu-6 cells. **B**, Time course of degradation of HiBit-CRAF by 1 µM ACBI-8451 in 293 cells. **C**, Degradation of endogenous GSPT1 in the indicated cell lines and of HiBit-GSPT1 in 293 cells. **D**, Degradation of endogenous CRAF and GSPT1 by CC-885 in SW620 cells at different time points. **E**, Time-course of degradation of HiBit-CRAF and HiBit-CRAF S259A by CC-885 in 293 cells. **F**, Degradation of HiBit fusions in 293 cells in the presence of cycloheximide (CHX). Blockade in protein translation negates any effects of the PROTAC on CRAF levels. **G**, Degradation of endogenous CRAF in 293 cells. Expression of the non-degradable GSPT1 G575N prevents effects on CRAF levels. For all dose response curves, treatment time was 18 hours unless indicated otherwise.

GSPT1 is a translation termination factor that has been shown to be degraded by the action of several molecular glues targeting cereblon ^23^, including PROTACs containing cereblon-binding moieties ^24,25^. Loss of GSPT1 causes a reduction in the rate of protein synthesis ^26^. Several of our hits, including ACBI-8451 (Figure 4C), were found to potently degrade both endogenous GSPT1 and its HiBit fusion, with degradation of both initiating within less than an hour of treatment.

A block in synthesis results in a reduction of all proteins according to their natural degradation rate. Since HiBit-CRAF S259A is a short-lived protein, we wondered if the observed degradation could be secondary to a reduction in protein synthesis caused by loss of GSPT1. To investigate this, we used CC-885, a known molecular glue that causes cereblon-dependent degradation of GSPT1. Degradation of GSPT1 by CC-885 caused a reduction in HiBit-CRAF S259A levels, with CRAF loss occurring with a time delay relative to GSPT1 degradation (Figure 4D, E), demonstrating that loss of GSPT1 can affect CRAF levels. We hypothesized that this was due to a reduction in protein translation caused by GSPT1 loss. If this assumption were correct, then in the absence of protein translation, ACBI-8451 would not further reduce CRAF levels. Indeed, when protein translation was blocked by cycloheximide, ACBI-8451 did not have an additional effect on CRAF levels, implying that all effects on CRAF protein levels are due to effects on protein synthesis, not degradation (Figure 4F). To directly determine if the degradation of GSPT1 is the only cause of the observed reduction in HiBit-CRAF S259A levels, we expressed the non-degradable GSPT1 mutant G575N^23^ in the cells (Figure 4G). This blocked degradation of HiBit-CRAF S259A by ACBI-8451, further demonstrating that it is a consequence of GSPT1 degradation.

If the decrease in CRAF levels observed after GSPT1 degradation is due to reduced protein synthesis, one would expect to also observe a general drop in the levels of other short-lived proteins. MYC, MCL1, MDM2 and ID2 were selected as proteins known to have a short half-life in 293 cells ^27^. We compared the effects of the known GSPT1 degrader CC-885 and ACBI-8451 to cycloheximide in A427 cells. As can be seen in Figure 5A, the effects are remarkably similar, with short-lived proteins such as MYC and MCL1 being reduced to lower levels than long-lived proteins such as SMARCA2. Note that CRAF is relatively short-lived in A427 cells (Figure 1A), and therefore relatively strongly affected.

**Figure 5.**
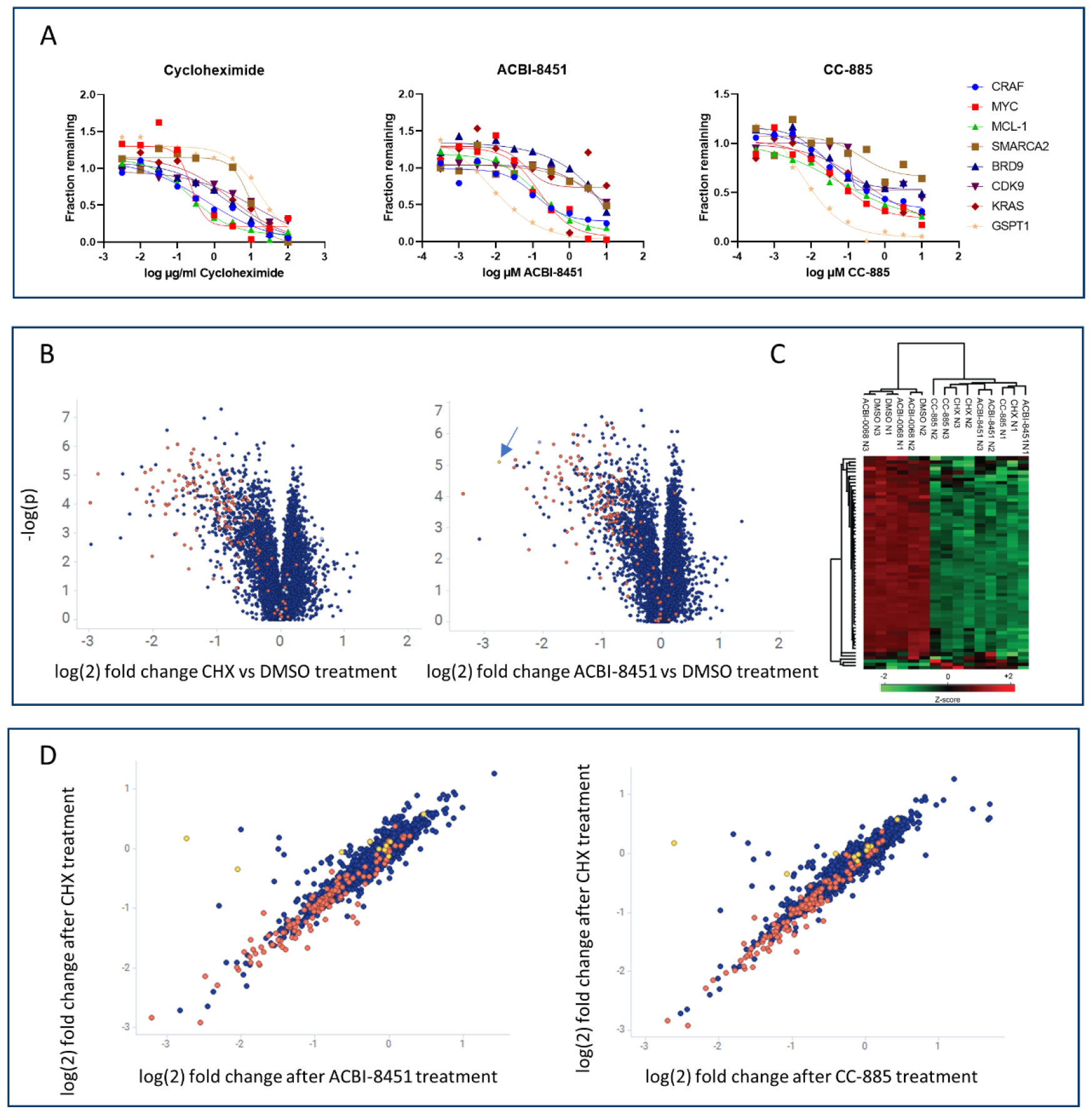
Degradation of GSPT1 phenocopies the effects of cycloheximide. **A**, Effect of the indicated compounds on protein levels after 18 hours of treatment in A427 cells. **B**, Levels of short–lived proteins are systematically lowered upon treatment by cycloheximide or ACBI-8451. HCT116 cells were treated overnight before protein abundance was assessed by mass spectrometry. Proteins were classified as short-lived (T ½ < 8 hours, red) and not short-lived (T ½ > 8 hours, blue) according to ^27^. Known neosubstrates of cereblon are indicated in yellow, GSPT1 is indicated by an arrow. **C**. Hierarchical clustering of short-lived proteins in the proteomic dataset with a significant difference in treated vs untreated samples, FDR < 0.05 (ANOVA). Green and red represent down-regulated and up-regulated levels of the proteins. ACBI-0068 is a non-cereblon-binding variant of ACBI-8451. **D**, Effects of cycloheximide and ACBI-8451 or CC-885 on changes in the levels of proteins are indistinguishable. Log(2) fold changes in protein abundance induced by the indicated compounds are plotted. Treatment of cells and coloring as in B.

In a proteome-wide study of the effects of one of our GSPT1 degrading compounds, this picture was further corroborated. After 24 hours of treatment with either cycloheximide or ACBI-8451, the abundance of many proteins with a half-life below 8 hours ^27^ was reduced (Figure 5B; known cereblon neosubstrates indicated in yellow; Table S1). Hierarchical clustering of samples indicated a high similarity of the effects of cycloheximide and ACBI-8451 treatment (Figure 5C). Interestingly, the effects of GSPT1 degradation and cycloheximide treatment were almost perfectly correlated (Figure 5D). This demonstrates that GSPT1 degradation has a very similar effect on protein abundance as cycloheximide. Both treatments reduce the levels of short-lived proteins by stalling protein synthesis. In summary, these data spell out the obvious expectation that a drop in protein synthesis leads to a relative reduction in the steady-state levels of short-lived proteins. This allows agents that stall protein synthesis to appear as agents that enhance the degradation of short-lived proteins such as MYC or MDM2, while all they do is lower the rate of protein synthesis. We conclude that an efficient GSPT1-degrading glue or PROTAC will always appear to be a degrader of most, if not all, short-lived proteins.

### Cytotoxic agents can cause a drop in short-lived proteins

After removing all GSPT1-degrading compounds, we were left with a small collection of hit compounds that appeared as non-GSPT1-driven degraders of HiBit-CRAF S259A. These compounds persisted in their apparent degradation of HiBit-CRAF S259A when the non-degradable GSPT1 G575N was co-expressed, as exemplified for ACBI-5767 in Figure 6A. However, these compounds also displayed a lag in time-resolved degradation measurements of CRAF (Figure 6B) as well as a drop in the levels of several short-lived proteins (Figure 6C). These compounds were equally active on versions of CRAF bearing mutations predicted to prevent compound binding (Fig. S1). We noted that these compounds, while inconspicuous after 24 hours of treatment, reduced cell viability after three days of treatment. We questioned whether cytotoxicity could be more generally associated with a reduction of the levels of short-lived proteins. Indeed, several cytotoxic agents caused a robust reduction in the levels of MDM2 and MCL1 (half-lives of 0,7 and 2,5 hours respectively ^27^, Figure 6D).

**Figure 6.**
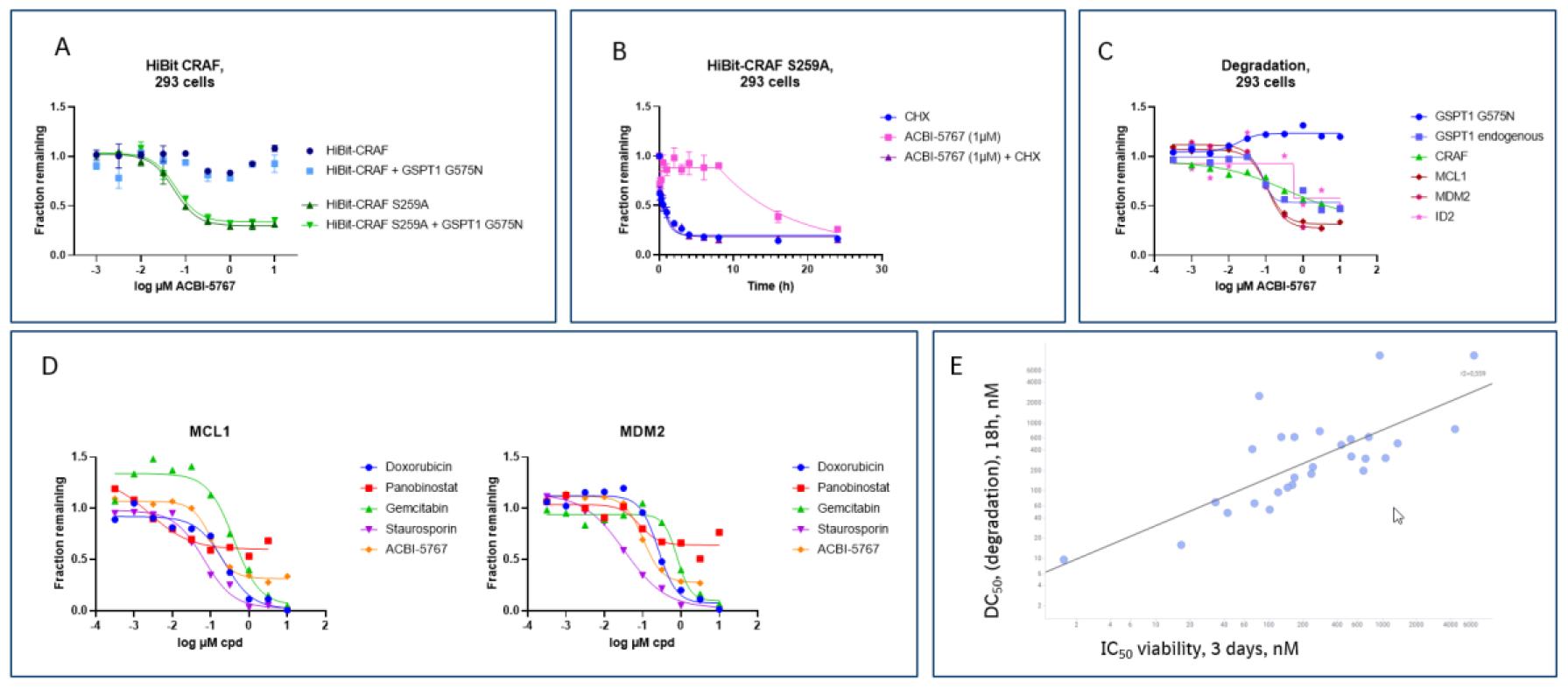
Deceptive degradation caused by cytotoxicity. **A**, The effects of ACBI-5767 are independent of GSPT1 degradation. Protein levels were measured using the HiBit assay after 18 hours of compound treatment in cell lines expressing HiBit-CRAF alone or HiBit-CRAF plus GSPT1 G575N. **B**, Time course of degradation of HiBit-CRAF S259A in 293 cells. **C**, Effect of ACBI-5767 on endogenous proteins after 18 hours of treatment. Protein levels were determined by capillary electrophoresis. **D**, Effect of cytotoxic compounds on endogenous proteins in 293 cells. Protein levels were determined by capillary electrophoresis. **E**, IC_50_ values as determined in viability assays are compared to DC_50_ values of HiBit-CRAF S259A in 293 cells expressing GSPT1 G575N. Axis labels are nanomolar.

The apparent degrading activities correlated significantly with cytotoxicity after three days of treatment, implying that one could be the reason for the other (Figure 6E). Given that endogenous CRAF is only partially degraded, and degradation of the exogenously expressed HiBit-CRAF S259A is unlikely to cause cytotoxicity, we are led to assume that cytotoxic properties of the compounds caused a drop in the synthesis rates of HiBit-CRAF S259A, leading to lower levels. We do not know why these compounds are cytotoxic, or if the drop in synthesis is related to reductions in transcription or translation. However, we can conclude that the effects are unrelated to GSPT1 degradation since all the assays were done in the presence of degradation resistant GSPT1 G575N. Given the limited number of compounds tested, we cannot conclude that all cytotoxic effects will necessarily reduce the levels of short-lived proteins.

### Competition experiments can differentiate PROTAC activity from artifactual effects

Most of the described artifacts cannot be captured by the typical controls of inhibiting proteasome activity or neddylation since a blockade in the natural ubiquitination and degradation process may equally affect the natural degradation pathways of these proteins. To exemplify this, we used blockage of transcription via CDK9 inhibition ^28^ to indirectly lower the levels of short-lived proteins in a dose-dependent manner. Figure 7A shows the dose dependent effect of transcriptional inhibition on protein levels, which is maximal for short-lived proteins such as MDM2 and MCL1, and not apparent for long-lived proteins such as SMARCA2 and GSPT1. As can be seen in Figure 7B, the classical controls for PROTAC activity, i.e., proteasome inhibition via MG132 or neddylation inhibition, can prevent the observed drop in protein levels for MDM2 and in part for MCL1, falsely qualifying these indirect effects as bona fide targeted degradation events. The same controls would falsely qualify the GSPT1 degrader ACBI-8451 as a bona fide degrader of CRAF (Figure 7C).

**Figure 7.**
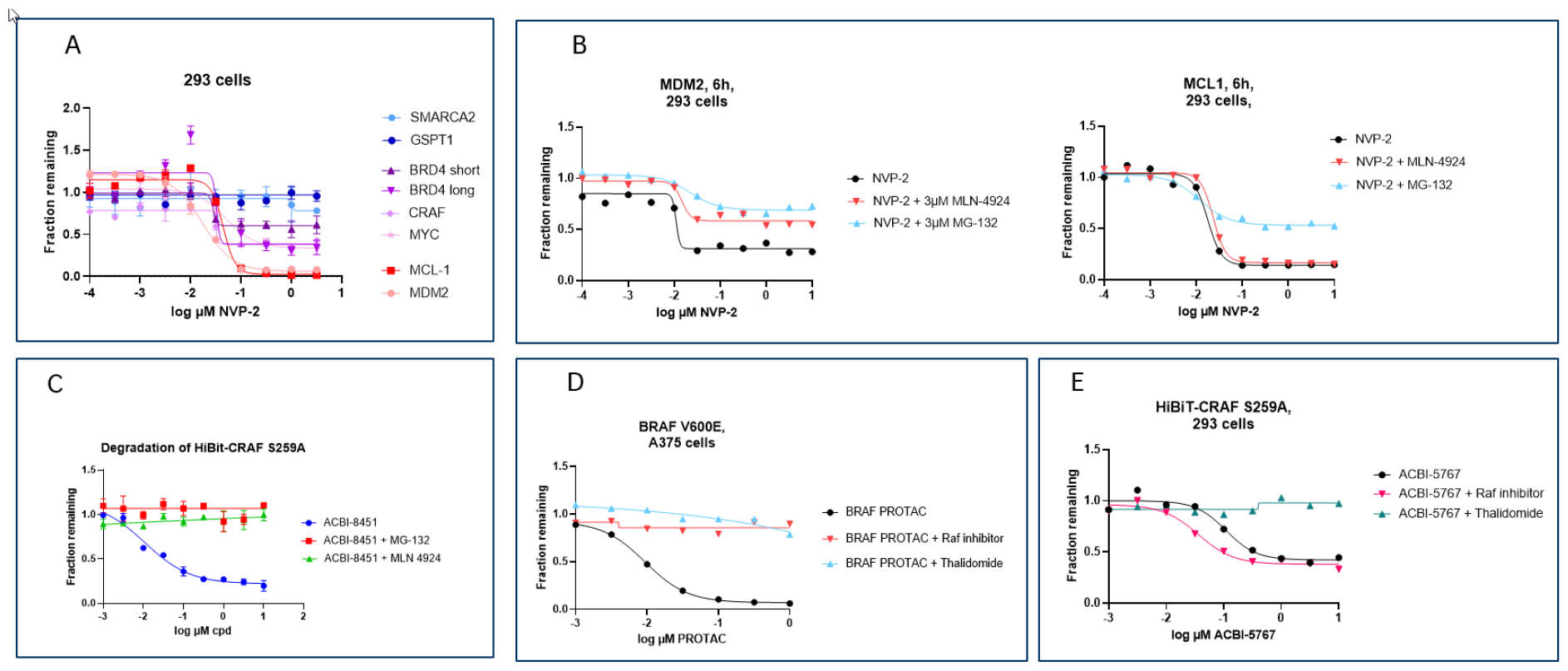
Controls to test for deceptive degradation. **A**, Effect of the transcriptional inhibitor NVP-2 on the levels of the indicated proteins, 18h. **B**, The effects of the transcriptional inhibitor NVP2 on MDM2 and MCL1 levels can in part be prevented by inhibitors of neddylation (MLN-4924) and the proteasome (MG-132). **C**, The effects of ACBI-8451 on HiBit-CRAF S259A can be prevented by inhibitors of neddylation (MLN 4924) or of the proteasome (MG-132). **D**, Degradation of BRAF V600E is efficiently prevented by competition with a BRAF binder or a cereblon binder. **E**. Degradation and proliferation assays of HiBit-CRAF-S259A in the presence or absence of compounds competing for binding to the target protein CRAF (ACBI-2370, a RAF inhibitor) or cereblon (4-OH-thalidomide). Note the left-shift in the presence of the RAF inhibitor.

In our case, competition experiments with RAF binding compounds proved to be distinctive controls: For a true PROTAC, competition for substrate or ligase binding is expected to prevent degradation. This can be exemplified here for a cereblon-based BRAF V600E PROTAC (WO2022261250A1) (Figure 7D).

In contrast, for the artifactual PROTACs, inclusion of a RAF-binding compound in the degradation assay did not prevent the observed reduction in HiBit-CRAF-S259A. On the contrary, the effects became more potent, presumably by adding to the cytotoxic effects (Figure 7E). Competition with 4-hydroxy thalidomide for CRBN binding equally blocks degradation. This shows that binding to cereblon is required for the artifactual effects to occur. Both observations suggest that the molecules are acting as either a molecular glue or as a PROTAC inducing degradation on unknown cytotoxicity-inducing off-targets. We would like to conclude that for PROTACs, especially for those targeting short-lived proteins, several controls are necessary to confirm their mode of action. These include a detailed time course of the degradation, an analysis of the effects on other short-lived proteins, as well as competition experiments with a binder of the target protein.

## Discussion

Removal of target proteins, be it by targeted protein degradation or genetic means, often leads to more pronounced effects than the mere inhibition of the target, as in the case of BCL6 ^29^ and CRAF ^18,30,31^. The irreversibility of protein degradation results in pharmacodynamic effects that can exceed the temporal confines of the exposure curve of a drug, which in the case of proteins with a low rate of synthesis, i.e. long-lived proteins, can markedly lengthen the span of target suppression *in vivo*. This is not true for short-lived proteins, which have a high rate of synthesis and will reach effective levels of protein shortly after the drug levels drop below active concentrations. In addition, it is more challenging to attain effective degraders for short-lived proteins since their levels will only be significantly depleted if the PROTAC-caused degradation rate significantly exceeds their already fast natural decay rate. ACBI2, a degrader of SMARCA2 and SMARCA4, which are long-lived proteins with half-lives exceeding 24 hours, achieves potent degradation of over 90% ^32^. We show here that a HiBit-tagged version of SMARCA2 displayed a shorter half-life and was consequently only partially degraded by ACBI1. This is in line with a mathematical model of protein degradation that takes the half-life/synthesis rate of the protein, the concentration of the PROTAC and the treatment time into consideration. This model can not only be used to predict efficacy of a drug in *in vitro* or *in vivo* models, but also the degradation rate required to effectively deplete levels of proteins of different natural stabilities, which may be of value when judging the feasibility of a project. It must be added here that while the induced degradation of short-lived proteins is demanding, even proteins as short-lived as MDM2 or MYC have been successfully degraded by bona fide small molecule degraders ^33,34^.

Cereblon-based PROTACs have been shown to inadvertently degrade proteins such as CSKN1A and GSPT1 via a molecular glue effect ^23,35^. Loss of GSPT1 reduces the rate of protein synthesis ^26^. Our data, including a proteomic analysis, demonstrate that effects of GSPT1 on protein abundance are indistinguishable from the effects of cycloheximide. Thus, any short-lived protein will deceivingly appear as a target of a GSPT1 degrader, even though the effect may be entirely indirect. GSPT1 degrading molecular glues are presently in early clinical development. It will be interesting to see if such a general cytotoxic effect can be efficacious in tumor patients at tolerable doses.

In addition, several cytotoxic compounds can indirectly cause the disappearance of short-lived proteins even before a loss in viability is observed, as measured by reduction in cellular ATP levels. We speculate that this is caused by a reduction in the rate of transcription and/or protein synthesis that a dying cell can maintain. While we cannot extend this observation to any cytotoxic agent, it is remarkable that agents as different as staurosporin or doxorubicin can elicit this effect.

These observations call for careful control experiments when reporting compounds which are designed to degrade short-lived proteins. We would like to propose that these controls should include:

- A degradation time course to specifically test for a delay in protein degradation after addition of the degrader compound. While cellular uptake may account for a delay in the range of minutes, delays in the range of hours highlight artifactual apparent degradation.
- Competition with binders; both to the POI and to the E3 ligase. As shown in this paper, indirect effects are not negated by competition experiments. If feasible, mutations in the target that prevent interaction with the PROTAC can be used to a similar intent. However, even this control can be misleading if there is off-target degradation that requires both the E3 ligase-binding and the POI-binding part of the PROTAC, and if this off-target degradation indirectly affects the levels of the POI.
- An analysis of the effects of the presumed degrader on other short-lived proteins, such as MCL1, MYC or MDM2. Their degradation would point to a broader effect on many short-lived proteins. Agents affecting many short-lived proteins are likely to work by mechanisms slowing the rate of protein biosynthesis. This is best tested via unbiased mass spectrometry proteomics at multiple timepoints.
- A combination of cycloheximide treatment with the PROTAC. Upon blockade of protein synthesis, the natural degradation rate is observed. Upon addition of both the degrader and cycloheximide, the degradation rate must increase over the natural degradation rate observed upon cycloheximide treatment alone.

Many of the controls that are frequently used to verify a PROTAC’s mode of action will not distinguish artifacts from bona fide degraders. These include inhibitors of the proteasome or of neddylation. Also, knock-outs of the E3 ligase or the use of compounds that have their E3-ligase binding part altered will not necessarily be informative in these cases. If GSPT1 degradation or cytotoxicity requires E3 ligase binding, the artifactual degradation will disappear with these controls.

While we do not know how some of the PROTACs in our study exerted their cytotoxic effect, we have seen similar effects with other PROTACs based on both cereblon and IAP binders, as well as a range of non-related cytotoxic compounds. Thus, we believe that it is very likely that such substances will be encountered in a screen of hundreds or thousands of candidate compounds. We hope that the practice of the outlined control experiments will help separate bona fide degraders from compounds acting indirectly.

## Significance

The use of protein degrading agents bears the potential to greatly extend the experimental possibilities in biomedical research. However, the use of molecules directly stalling protein synthesis, or of cytotoxic compounds that indirectly affect the abundance of short-lived proteins, has the potential to mislead and result in incorrect interpretations. Defining mechanisms, such as GSPT1 degradation, that cause the artifactual classification of small molecules as protein degraders will help to focus on true protein degrading molecules, thus saving time and effort from being spent following non-productive routes of research.

## Supporting information

Supplement Vetma et al.

## Acknowledgements

We are grateful to Anita Lehner and Peggy Stolt-Bergner for critical discussions and providing reagents for the CRAF project, and to Peter Ettmayer for critical discussions.

## Author contributions

AS, AK and TG designed and executed biological experiments. JE did mathematical analysis, performed numerical simulations and wrote a section of the paper. NB contributed to scientific discussions and executed screening and counter-screening activities for the PROTAC libraries. TG contributed to the construction of PROTAC libraries. YW supported the generation of RAF mutants not binding to inhibitors. WF and CK contributed to supervision of the project and experiment design. VV designed and executed biological and proteomics experiments and oversaw biology efforts in the early stage of the project. VS supported GSPT1 and CRAF protein production and general structural biology on the project. ED and NT were involved in initial PROTAC design efforts. ED, NT and ABF contributed to the synthesis of the compounds described within this manuscript. ABF and NC oversaw chemistry efforts on the project, and contributed to scientific discussions and experimental design. IP assisted in preparation of samples for proteomics and data analysis. KMcA contributed to supervision of the project and scientifc discussion. LCP designed and executed biological experiments and oversaw biology efforts in the later stage of the project. GK contributed to PROTAC design and synthesis. CW contributed to scientific discussions. TC and GD synthesised PROTAC libraries. PG oversaw chemistry efforts on PROTAC libraries and contributed to experimental design. AC supervised the project. MK supervised the project, designed experiments and wrote the paper.

## STAR Methods

### Chemical synthesis

Synthesis of compounds described in this paper and their intermediates are described in the Supplemental Information.

### Cell lines and culture

Cell lines were obtained through ATCC, verified for identity by satellite repeat analysis and tested for mycoplasma contamination. Cells were grown in the medium as specified by the supplier unless described otherwise and not used beyond passage 25.

### Proliferation assays

A total of 1,000 cells per well were seeded in 384-well plates. After overnight incubation, compounds were added to the cells at logarithmic dose series using the HP Digital Dispenser D300 (Tecan), normalizing for added DMSO. On the day of compound additions (=T0) or after the indicated time, cellular ATP content was measured using CellTiterGlo (Promega). Measurements were normalized to DMSO-treated cells to derive relative inhibition,

### Protein degradation assays

For capillary electrophoresis, 40,000 cells in 100 µl per well were seeded in a 96-well plate and incubated at 37 °C overnight. Compounds were added from DMSO stock solution using a Digital Dispenser D300 (Tecan), normalizing for added DMSO and cells were incubated at 37 °C for the indicated times. Medium was removed, cells washed with PBS and lyzed in 20 µl lysis buffer (1% Triton, 350 mM KCl, 10 mM TRIS pH 7.4, phosphatase-protease inhibitor cocktail (Thermo Scientific no. 1861281), 10 mM DTT, benzonase 0.5 µl ml™1 (Novagen no. 70746 10KU, 25 U per µl)) for 30 min on ice before insoluble debris was pelleted by centrifugation. Protein levels were determined on a JESS capillary electrophoresis instrument (Proteinsimple) using MCL1 (CellSignaling #5453, 1:50), MDM2 (CellSignaling #86934, 1:50), MYC (CellSignaling #5605S, 1:50), CRAF (CellSignaling 9422S, 1:25), SMARCA2 (Sigma HPA029981, 1:25), SMARCA4 (CellSignaling #49360, 1:25), GSPT1 (Abcam #ab49878, 1:50), BRD9 (Bethyl #A303-781A-1, 1:50), KRAS (LSBio #LS-C175665, 1:100), CDK9 (CellSignaling #2316, 1:50) and GAPDH (Abcam ab8245, 1:1000).

For HiBit assays, the following sequence coding for the HiBit peptide as well as an 8 amino acid linker was fused to the 5’ coding sequence of the indicated genes: ATGGTGAGCGGCTGGCGGCTGTTCAAGAAGATCTCCGGTGGTGGTGGTAGTGGTGGTGGTAGT and the construct was retrovirally expressed in 293 cells. 2500 cells per well of a 384-well plate were seeded before compound addition via the HP Digital Dispenser. After incubation for the indicated time, HiBit levels were detected using the Nano-Glo® HiBiT lytic detection system from Promega (#N3040).

To determine the half-life of proteins, cycloheximide was added to the cells (100 µg/ml, SIGMA). CDK9 inhibition was by NVP2 (MedChemExpress HY-12214A). CC-885 was obtained at MedChemExpress (HY-101488), MLN-4924 at ActiveBiochem (A-1139) and MG-32 at Merck (M7449).

### Proteomics

#### Sample preparation

293 cells in DMEM (Gibco) were seeded at 3 × 10^5^ cells on a 100 mm plate 24 h before treatment. Cells were treated in triplicate by addition of compounds ACBI-8451, CC-885, and ACBI-0068, a non-cereblon binding control version of ACBI-8451 at 1 µM and cycloheximide at 100 µg/mL. After 24 h, the cells were washed twice with 10 mL of cold PBS and lysed in 500 µL of 100 mM TEAB with 6% (w/v) SDS. The lysate was pulse sonicated briefly and then centrifuged at 15,000 × *g* for 10 min. Samples were quantified using a microBCA protein assay kit (Thermo Fisher Scientific). 300 µg of each sample was reduced with DTT, alkalised with iodoacetamide and double-digested with trypsin using the modified S-TRAP mini (ProtiFi) protocol. Peptide quantification was done using Pierce™ Quantitative Fluorometric Peptide Assay and equal amount from each sample was labelled using TMTpro™ 16plex Label Reagent Set Set (Thermo Fisher Scientific) as per the manufacturer’s instructions. The samples were then pooled and desalted using a 7 mm, 3 mL C18 SPE cartridge column (Empore, 3M). The pooled and desalted sample was fractionated using high pH reverse-phase chromatography on an XBridge peptide BEH column (130 Å, 3.5 μm, 2.1 × 150 mm, Waters) on an Ultimate 3000 HPLC system (Thermo Scientific/Dionex). Buffers A (10 mM ammonium formate in water, pH 9) and B (10 mM ammonium formate in 90% acetonitrile, pH 9) were used over a linear gradient of 2% to 100% buffer B over 80 min at a flow rate of 200 μL/min. 80 fractions were collected using a WPS-3000 FC auto-sampler (Thermo Scientific) before concatenation into 21 fractions based on the UV signal of each fraction. All the fractions were dried in a Genevac EZ-2 concentrator and resuspended in 1% formic acid for MS analysis.

### LC-MS/MS analysis

The fractions were analyzed sequentially on a Q Exactive HF Hybrid Quadrupole-Orbitrap Mass Spectrometer (Thermo Scientific) coupled to an Dionex Ultimate 3000 RS (Thermo Scientific). Buffers A (0.1% formic acid in water) and B (0.1% formic acid in 80% acetonitrile) were used over a linear gradient from 5% to 35% buffer B over 125 min and then from 35% buffer B to 98% buffer B in 2 min at a constant flow rate of 300 nL/min. The column temperature was 50 °C. The mass spectrometer was operated in data dependent mode with a single MS survey scan from 335-1600 *m/z* followed by 15 sequential *m/z* dependent MS2 scans. The 15 most intense precursor ions were sequentially fragmented by higher energy collision dissociation (HCD). The MS1 isolation window was set to 0.7 *m/z* and the resolution set at 120,000. MS2 resolution was set at 60,000. The AGC targets for MS1 and MS2 were set at 3×10^6^ ions and 1×10^5^ ions, respectively. The normalized collision energy was set at 32%. The maximum ion injection times for MS1 and MS2 were set at 50 ms and 200 ms respectively. The mass accuracy was checked before the initiation of sample analysis.

### Peptide and protein identification

The raw MS data files for all 20 fractions were merged and searched against the Uniprot-sprot-Human-Canonical database by Maxquant software 1.4.16 for protein identification and TMT reporter ion quantitation. The Maxquant parameters were set as follows: enzyme used Trypsin/P; maximum number of missed cleavages equal to two; precursor mass tolerance equal to 10 p.p.m.; fragment mass tolerance equal to 20 p.p.m.; variable modifications: oxidation (M), dioxidation (MW), acetyl (N-term), deamidation (NQ), Gln -> pyro-Glu (Q N-term); fixed modifications: carbamidomethyl (C). The data was filtered by applying a 1% false discovery rate followed by exclusion of proteins with less than two unique peptides. Quantified proteins were filtered if the absolute fold-change difference between the three DMSO replicates was ≥ 1.5. The mass spectrometry proteomics data have been deposited to the ProteomeXchange Consortium via the PRIDE ^36^ partner repository with the dataset identifier PXD047934.

