## Supplement Vetma et al. for "Confounding factors in targeted degradation of short-lived proteins"

**Summary**
Targeted protein degradation has recently emerged as a novel option in drug discovery. Natural protein half-life is expected to affect the efficacy of degrading agents, but to what extent it influences target protein degradation has not been systematically explored. Using mathematical modelling of protein degradation, we demonstrate that the natural half-life of a target protein has a dramatic effect on the level of protein degradation induced by a degrader agent which can pose significant hurdles to screening efforts. Moreover, we show that upon screening for degraders of short-lived proteins, agents that stall protein synthesis, such as GSPT1 degraders and generally cytotoxic compounds, deceptively appear as protein degrading agents. This is exemplified by the disappearance of short-lived proteins such as MCL1 and MDM2 upon GSPT1 degradation and upon treatment with cytotoxic agents such as doxorubicin. These findings have implications for target selection as well as for the type of control experiments required to conclude that a novel agent works as a bona fide targeted protein degrader.

A

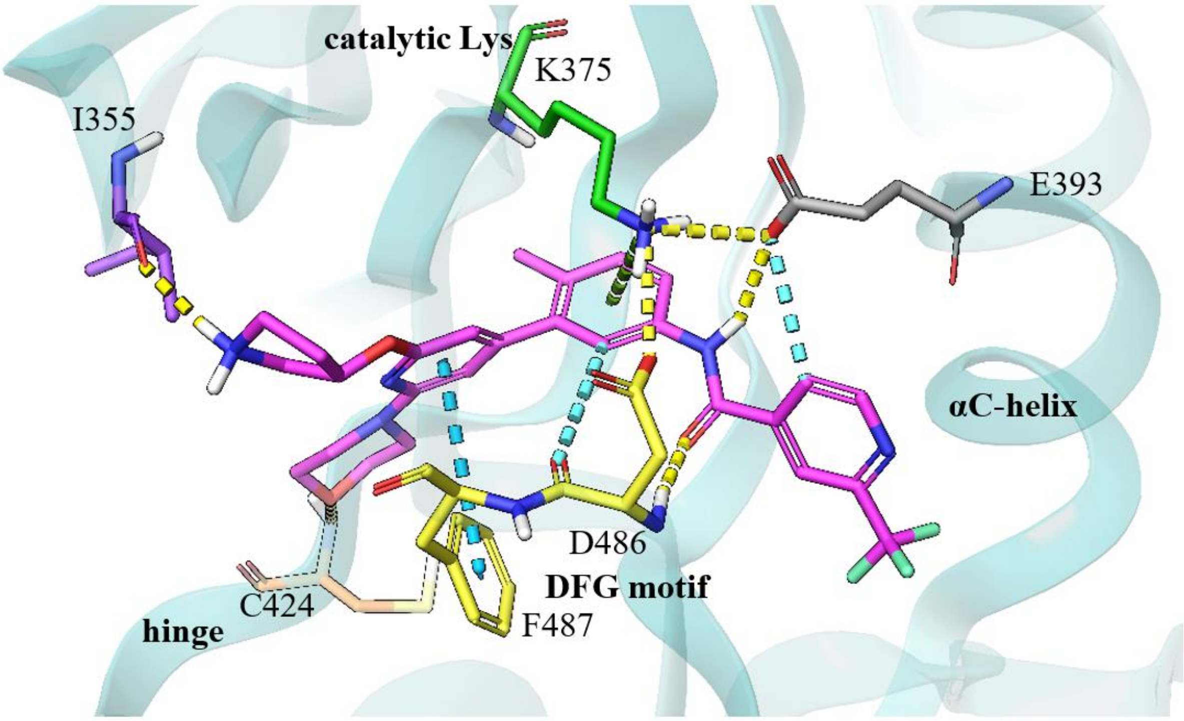


B

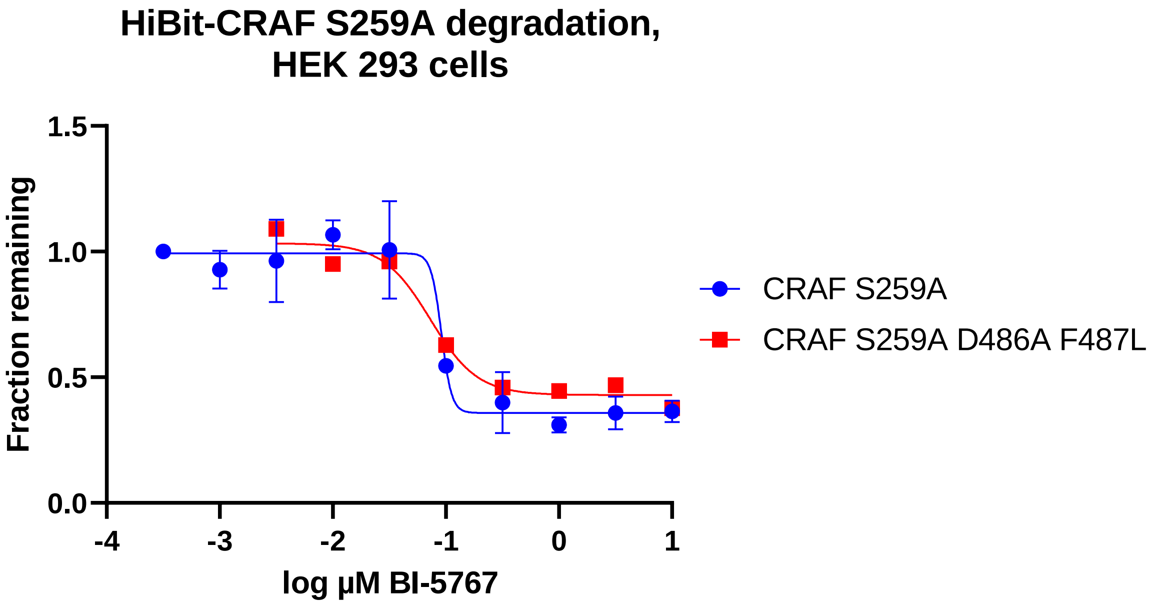


Figure S1: **A**, Rationale for CRAF mutants. To accommodate the CRAF binder of the PROTAC ACBI-5767, a homology model of CRAF with an accessible back pocket was built based on a public BRAF structure with LXH254 (PDB ID: 8F7P). The secondary structure of CRAF is represented with cyan ribbons. The CRAF binder of ACBI-5767 and relevant binding site residues are shown as sticks and colored by atom type, with magenta carbons for the binder. Residue carbon atoms are colored according to their locations in the binding site: DFG motif in yellow (only D486 and F487 are displayed), hinge in orange, solvent-exposed in violet, catalytic lysine in green, and αC-helix glutamic acid in grey. H-bonds and salt bridges are shown as yellow dashed lines, π-π stacking and aromatic H-bond interactions as cyan dashed lines, and a cation-pi interaction as green dashed line. On the left, the morpholine forms a weak H-bond with the backbone of C424 at the hinge, and the pyrrolidinium forms an H-bond to the solvent-exposed I355. The pyridyl at the center is stabilized by a π-π stacking with F487 of the DFG motif. The methylphenyl forms a cation-π interaction with the catalytic K375 and an aromatic H-bond to D486 of the DFG motif. K375 is also held in place by salt bridges with D486 of the DFG motif and E393 of the αC-helix. The binder amide undergoes a salt bridge with E393 and an H-bond with D486. The trifluoromethylpyridine makes an aromatic H-bond to E393. LXH254 in the template BRAF crystal structure forms slightly weaker, but similar interactions. The pyridyl ring makes a π-π stacking with F595 of the DFG motif and the methylphenyl ring in the center of the structure makes a cation-π interaction with the catalytic K483. The amide is fixed by D594 of the DFG motif and E501 of the αC-helix. In both structures, the CRAF model with the binder of PROTAC ACBI-5767 and the BRAF X-ray structure with LXH254, the pyridyl and the methylphenyl feature extensive hydrophobic interactions with the DFG motif. The binding mode of the binder of the PROTAC ACBI-5767 in the CRAF model shown is compatible with the exit vector of the fully decorated PROTAC. Based on the model, a CRAF D486A F487L mutant cannot bind the PROTAC ACBI-5767.

**B**, Degradation assay for HiBit-CRAF S259A D486A F487L. Since this mutant cannot bind to the PROTAC, the observed degradation must be an indirect effect. The HiBit-CRAF S259A background was chosen since it has a stronger response to ACBI-5767 than the wild type HiBit-CRAF.

Table S1. Cereblon neo-substrate list compiled from literature [1–5]. Proteins identified in our proteomics dataset are marked "yes" and highlighted in yellow in the table and in Fig.5B. Other proteins were not identified in our proteomics dataset and are marked "not found".

| CRBN Neo-substrate | Uniprot ID | identified in proteomics dataset | CRBN Neo-substrate | Uniprot ID | identified in proteomics dataset |
|---|---|---|---|---|---|
| RAB28 | P51157 | yes | ZNF692 | Q9BU19 | not found |
| ZFP91 | Q96JP5 | yes | SALL4 | Q9UJQ4 | not found |
| DTWD1 | Q8N5C7 | yes | RNF166 | Q96A37 | not found |
| CSNK1A1 | P48729 | yes | FAM83F | Q8NEG4 | not found |
| GSPT1 | P15170 | yes | IKZF1 | Q13422 | not found |
| ZMYM2 | Q9UBW7 | yes | IKZF2 | Q9UKS7 | not found |
| ZNF143 | P52747 | yes | ZNF827 | Q17R98 | not found |
| WIZ | O95785 | yes | GZF1 | Q9H116 | not found |
| PATZ1 | Q9HBE1 | yes | ZBTB39 | O15060 | not found |
| ZNF787 | Q6DD87 | yes | ZNF653 | Q96CK0 | not found |
| ZNF98 | A6NK75 | not found | IKZF4 | Q9H2S9 | not found |
| ZBTB16 | Q05516 | not found | ZKSCAN5 | Q9Y2L8 | not found |
| IKZF3 | Q9UKT9 | not found | ZNF582 | Q96NG8 | not found |
| SALL3 | Q9BXA9 | not found | ZNF517 | Q6ZMY9 | not found |
| E4F1 | Q66K89 | not found | ZNF276 | Q8N554 | not found |
| ZNF654 | Q8IZM8 | not found | | | |

1. Furihata, H., Yamanaka, S., Honda, T., Miyauchi, Y., Asano, A., Shibata, N., Tanokura, M., Sawasaki, T., and Miyakawa, T. (2020). Structural bases of IMiD selectivity that emerges by 5-hydroxythalidomide. Nat. Commun. *11*, 4578. 10.1038/s41467-020-18488-4.

2. Yamanaka, S., Furihata, H., Yanagihara, Y., Taya, A., Nagasaka, T., Usui, M., Nagaoka, K., Shoya, Y., Nishino, K., Yoshida, S., et al. (2023). Lenalidomide derivatives and proteolysis-targeting chimeras for controlling neosubstrate degradation. Nat. Commun. *14*, 4683. 10.1038/s41467-023-40385-9.

3. Donovan, K.A., An, J., Nowak, R.P., Yuan, J.C., Fink, E.C., Berry, B.C., Ebert, B.L., and Fischer, E.S. (2018). Thalidomide promotes degradation of SALL4, a transcription factor implicated in Duane Radial Ray syndrome. eLife *7*, e38430. 10.7554/elife.38430.

4. Nguyen, T.M., Sreekanth, V., Deb, A., Kokkonda, P., Tiwari, P.K., Donovan, K.A., Shoba, V., Chaudhary, S.K., Mercer, J.A.M., Lai, S., et al. (2024). Proteolysis-targeting chimeras with reduced off-targets. Nat. Chem. *16*, 218–228. 10.1038/s41557-023-01379-8.

5. Sievers, Q.L., Petzold, G., Bunker, R.D., Renneville, A., Slabicki, M., Liddicoat, B.J., Abdulrahman, W., Mikkelsen, T., Ebert, B.L., and Thoma, N.H. (2018). Defining the human C2H2 zinc finger degrome targeted by thalidomide analogs through CRBN. Science *362*, eaat0572. 10.1126/science.aat0572.

Chemical synthesis

#### General information

All chemicals unless otherwise stated, were commercially available, at least 90% pure and used without further purification. Commercially available dry solvents were used. All reactions were carried out in oven- or flame dried glassware under nitrogen atmosphere. Normal phase TLC was carried out on pre-coated silica plates (Kieselgel 60 F254, BDH) with visualization via UV light (UV 254 and/or 365 nm) and/or basic potassium permanganate solution. Isolute® phase separator columns from Biotage were used. Flash column chromatography (FCC) was performed using either a Teledyne Isco Combiflash Rf or Rf200i or a Biotage Isolera One with prepacked Redisep RF Normal phase disposable Columns. Reverse phase chromatography was carried out on a Biotage Isolera One using SNAP-C18 Columns. NMR Spectra were recorded on a Bruker Ascend 400 MHz or 500 MHz as specified. Chemical shifts are quoted in ppm and referenced to the residual solvent signals: $^{1}$H NMR $\delta$ (ppm) = 7.26 ($CDCl_3$), $^{13}$C NMR $\delta$ (ppm) = 77.16 ($CDCl_3$); $^{1}$H NMR $\delta$ (ppm) = 5.32 ($CD_2Cl_2$), $^{13}$C NMR $\delta$ (ppm) = 53.84 ($CD_2Cl_2$), $^{1}$H NMR $\delta$ (ppm) = 2.50 (DMSO-$d_6$). Signal splitting patterns are described as singlet (s), doublet (d), triplet (t), quartet (q), multiplet (m), broad (br) or a combination thereof. Coupling constants ($J$) are measured in Hertz (Hz). High Resolution Mass Spectra (HRMS) were recorded on a Bruker microTOF. Other resolution MS and analytical HPLC traces were recorded on an Agilent Technologies 1200 series HPLC connected to an Agilent Technologies 6130 quadrupole LC/MS, connected to an Agilent diode array detector. The column used was a Waters XBridge column (50 mm × 2.1 mm, 3.5 µm particle size) and the compounds were eluted with a gradient 5−95% acetonitrile/water + 0.1% formic acid ("acidic method").
Preparative HPLC was performed on a Waters Preparative HPLC System with a Waters XBridge C18 column (100 mm x 19 mm; 5 µm particle size) and a gradient of 5–95% acetonitrile in water over 10 minutes, flow 25 mL/min, with 0.1% ammonia in the aqueous phase.
Intermediate **4** was synthesised as per the procedure described in Crew, A. P.; Hornberger, K. R.; Wang, J. Crews, C. M.; Jaime-Figueroa, S.; Dong, H.; Qian, Y.; Zimmerman, K. U.S. Patent 0129627 A1, 2020.
All compounds used in biological assays were >95% purity by LCMS analysis unless otherwise stated.

Abbreviations used: aq. for aqueous, DCE for 1,2-dichloroethane, DCM for dichloromethane, DIPEA for *N*,*N*-diisopropylethylamine, DMAP for N,N-dimethylpyridin-4-amine, DME for 1,2-Dimethoxyethane, DMF for *N*,*N*-dimethylformamide, DMSO for dimethylsulfoxide, equiv. for equivalents, EtOAc for ethyl acetate, EtOH for ethanol, HATU for 1-[bis(dimethylamino)methylene]-1*H*-1,2,3-triazolo[4,5-*b*]pyridinium 3-oxid hexafluorophosphate, HCl for hydrochloric acid, HPLC for high-performance liquid chromatography, LCMS for liquid chromatography mass spectrometry, MeCN for acetonitrile, MeI for methyl iodide, MeOD for methanol-$d_6$, MeOH for methanol, NaH for sodium hydride, NaOH for sodium hydroxide, NMP for *N*-methyl-2-pyrrolidone, sat. for saturated, TFA for trifluoroacetic acid, TsCl for tosyl chloride.

#### Synthesis of N-(6'-((1-(3-(2-((2-(2,6-dioxopiperidin-3-yl)-1-oxoisoindolin-5-yl)oxy)ethoxy)propanoyl)piperidin-4-yl)oxy)-2-methyl-5'-morpholino-[3,3'-bipyridin]-5-yl)-3-(trifluoromethyl)benzamide (ACBI-8451)

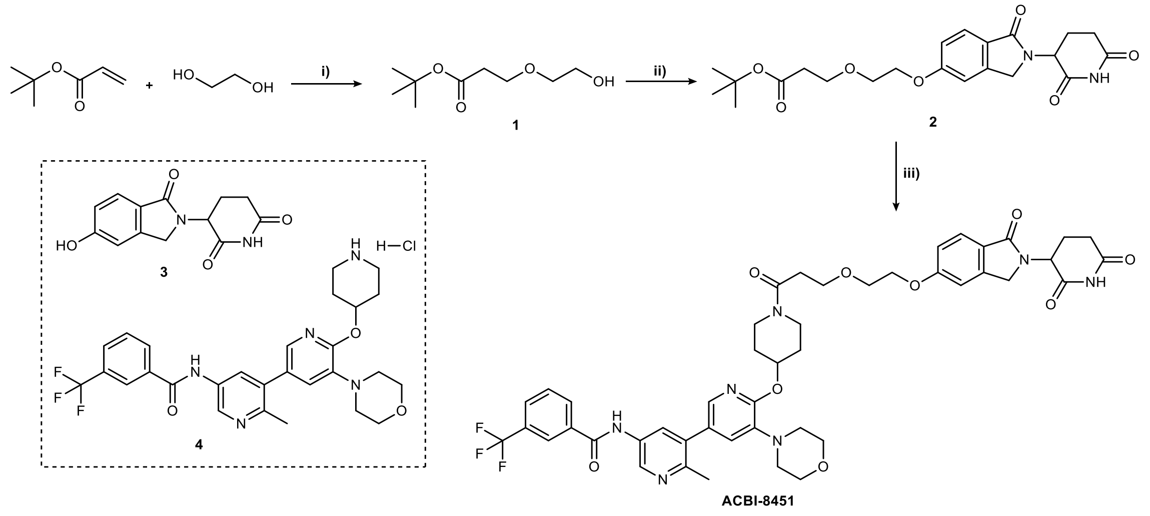


**Synthetic route to compound ACBI-8451.** Reagents and conditions: i) TRITON B (40% in $H_2O$), MeCN, rt, 18 h; ii) **3**, DIAD, $PPh_3$, THF, NMP, 50°C, 16 h; iii) a.TFA, $CH_2Cl_2$, rt, 18 h; then b. **4**, HATU, DIPEA, $CH_2Cl_2$, rt, 10 min.

##### *tert*-butyl 3-(2-hydroxyethoxy)propanoate, 1

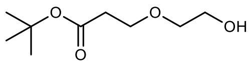

TRITON B, 40% in water (1.0 mL, 3.0 mmol) was added to a solution of ethane-1,2-diol (17.2 mL, 308 mmol) and *tert*-butyl acrylate (9.2 mL, 63 mmol) in MeCN (40 mL). The resulting reaction mixture was stirred at room temperature for 18 h, then concentrated under reduced pressure. The remaining residue was diluted with $CH_2Cl_2$ and $H_2O$, then passed through a phase separator. The organic phase was concentrated under reduced pressure. The remaining residue was purified by column chromatography on silica gel (0-100% EtOAc in Heptane) to yield **1** as a colourless oil (3.55 g, 30% yield).

$^1H$ NMR (400 MHz, $CDCl_3$) $\delta$ (ppm) = 3.71 – 3.60 (m, 4H), 3.50 (dd, $J$ = 5.2, 3.9 Hz, 2H), 2.43 (t, $J$ = 6.1 Hz, 2H), 2.31 (t, $J$ = 6.3 Hz, 1H), 1.39 (s, 9H).

##### *tert*-butyl 3-(2-((2-(2,6-dioxopiperidin-3-yl)-1-oxoisoindolin-5-yl)oxy)ethoxy)propanoate, 2

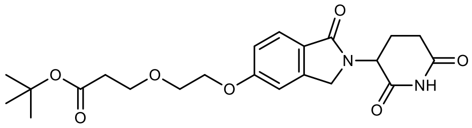

$PPh_3$ (378 mg, 1.44 mmol) was added to a 50°C solution of 3-(5-hydroxy-1-oxoisoindolin-2-yl)piperidine-2,6-dione, **3** (300 mg, 1.15 mmol) and *tert*-butyl 3-(2-hydroxyethoxy)propanoate, **1** (548 mg, 2.88 mmol) in THF (14.0 mL, 175 mmol) and N-Methyl-2-pyrrolidinone (NMP) (3.0 mL). The resultant mixture was stirred for 5 minutes, before addition of Diisopropyl azodicarboxylate (454 µl, 2.31 mmol). The mixture was stirred for 16 h at 50°C, and analysis by LCMS showed reaction completion. The reaction was quenched by addition of $H_2O$, then concentrated under reduced pressure. The remaining residue in NMP was purified by reverse phase column chromatography under acidic conditions (5-95% $CH_3CN$ in 0.1% aq. $HCO_2H$) to give the desired product, contaminated with a $PPh_3O$ impurity. Further purification by normal phase chromatography (0-4% MeOH in DCM) gave **2** as an amorphous yellow solid (143 mg, 29% yield).

$^{1}$H NMR (400 MHz, $CDCl_3$) $\delta$ (ppm) = 8.76 (s, 1H), 7.66 (d, *J* = 8.5 Hz, 1H), 6.90 (dd, *J* = 8.5, 2.2 Hz, 1H), 6.85 (d, *J* = 2.2 Hz, 1H), 5.08 (dd, *J* = 13.2, 5.2 Hz, 1H), 4.30 (d, *J* = 16.1 Hz, 1H), 4.16 (d, *J* = 16.1 Hz, 1H), 4.07 (dd, *J* = 5.7, 3.9 Hz, 2H), 3.79 – 3.61 (m, 4H), 2.80 – 2.62 (m, 2H), 2.43 (t, *J* = 6.4 Hz, 2H), 2.20 (tdd, *J* = 13.0, 11.1, 6.8 Hz, 1H), 2.10 – 1.99 (m, 1H), 1.35 (s, 9H).

**Synthesis of N-(6'-((1-(3-(2-((2-(2,6-dioxopiperidin-3-yl)-1-oxoisoindolin-5-yl)oxy)ethoxy)propanoyl)piperidin-4-yl)oxy)-2-methyl-5'-morpholino-[3,3'-bipyridin]-5-yl)-3-(trifluoromethyl)benzamide (ACBI-8451)**

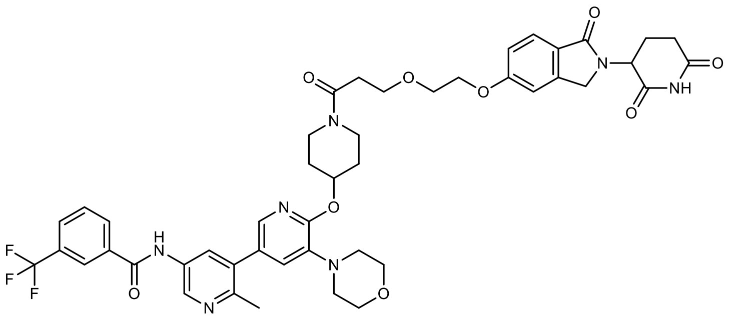


TFA (1.0 mL) was added to a solution of *tert*-butyl 3-(2-((2-(2,6-dioxopiperidin-3-yl)-1-oxoisoindolin-5-yl)oxy)ethoxy)propanoate, **2** (60 mg, 0.14 mmol) in DCM (1.0 mL) at 0°C. The reaction mixture was stirred at room temperature for 18 h. LCMS showed full conversion of the *tert*-butyl ester to the corresponding carboxylic acid. The solvents were removed by aspiration under nitrogen and azeotroped twice with toluene. A mixture of the crude acid, *N*-(2-methyl-5'-morpholino-6'-(piperidin-4-yloxy)-[3,3'-bipyridin]-5-yl)-3-(trifluoromethyl)benzamide hydrochloride, **4** (80 mg, 0.14 mmol), HATU (79 mg, 0.21 mmol) and DIPEA (113 µl, 0.69 mmol) in DCM (1.0 mL) was stirred at room temperature for 10 minutes. LCMS analysis showed majority conversion to desired product. Purification by normal phase column chromatography (0-8% MeOH in DCM), followed by preparative HPLC purification under acidic conditions (5-95% $CH_3CN$ in 0.1% aq. HCOOH) gave **ACBI-8451** as an amorphous white solid (81 mg, 62% yield).
$^{1}$H NMR (400 MHz, $CDCl_3$) $\delta$ (ppm) = 8.80 (d, *J* = 19.0 Hz, 1H), 8.69 (dd, *J* = 16.1, 2.5 Hz, 1H), 8.37 (br s, 1H), 8.16 – 8.13 (m, 1H), 8.09 – 8.05 (m, 1H), 8.02 (dd, *J* = 6.5, 2.5 Hz, 1H), 7.71 (d, *J* = 7.8 Hz, 1H), 7.64 – 7.61 (m, 1H), 7.59 – 7.50 (m, 2H), 6.96 – 6.94 (m, 1H), 6.87 – 6.83 (m, 2H), 5.33 – 5.27 (m, 1H), 4.97 (dd, *J* = 13.2, 5.1 Hz, 1H), 4.31 (d, *J* = 16.0 Hz, 1H), 4.17 (d, *J* = 16.0 Hz, 1H), 4.08 – 4.05 (m, 2H), 3.82 – 3.71 (m, 9H), 3.66 – 3.59 (m, 1H), 3.58 – 3.51 (m, 1H) 3.47 – 3.37 (m, 1H), 3.03 – 2.98 (m, 4H), 2.75 (ddd, *J* = 17.7, 4.6, 2.4 Hz, 1H), 2.68 – 2.52 (m, 4H), 2.40 (s, 3H), 2.28 – 2.13 (m, 1H), 2.09 – 1.96 (m, 1H), 1.93 (dd, *J* = 13.1, 4.2 Hz, 1H), 1.77 (s, 2H). HRMS (ESI) for $C_{46}H_{48}F_3N_7O_9$ $[M+H]^+$ calculated 900.3538, obtained 900.3545.

#### Synthesis of N-(2-methyl-6'-((1-(3-(2-((2-(1-methyl-2,6-dioxopiperidin-3-yl)-1-oxoisoindolin-5-yl)oxy)ethoxy)propanoyl)piperidin-4-yl)oxy)-5'-morpholino-[3,3'-bipyridin]-5-yl)-3-(trifluoromethyl)benzamide, (ACBI-0068)

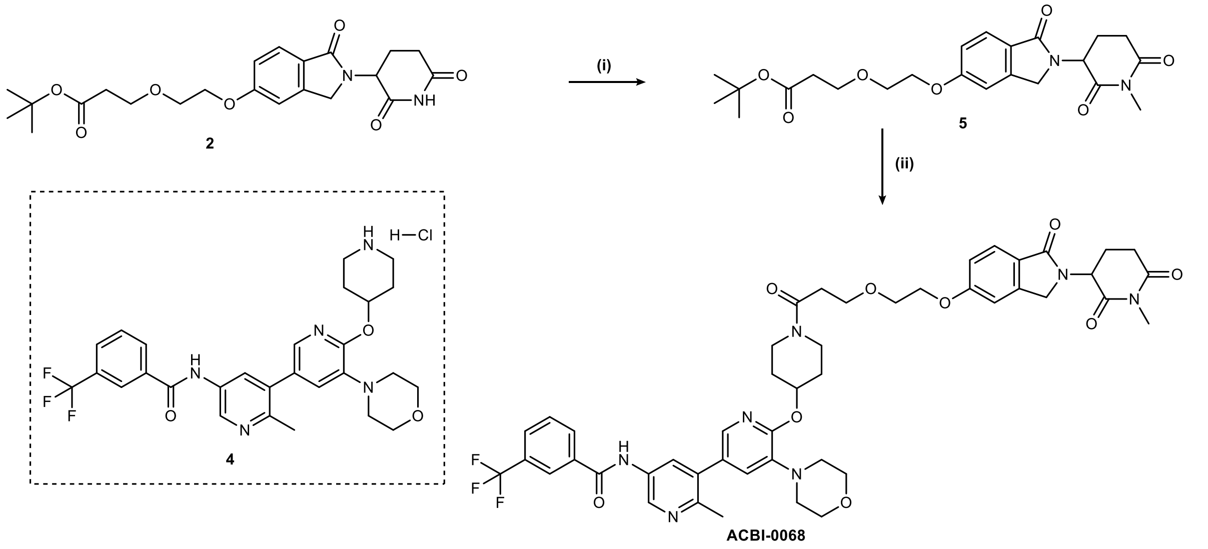


**Synthetic route to compound ACBI-0068.** Reagents and conditions: i) MeI, $K_2CO_3$, DMF, rt, 18 h; ii) a.TFA, $CH_2Cl_2$, rt, 18 h; then b. **4**, HATU, DIPEA, $CH_2Cl_2$, rt, 10 min.

#### *tert*-butyl 3-(2-((2-(1-methyl-2,6-dioxopiperidin-3-yl)-1-oxoisoindolin-5-yl)oxy)ethoxy)propanoate, 5

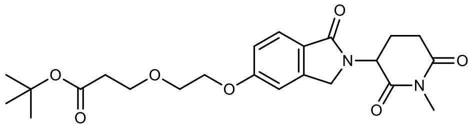

*tert*-Butyl 3-(2-((2-(2,6-dioxopiperidin-3-yl)-1-oxoisoindolin-5-yl)oxy)ethoxy)propanoate, **2** (29 mg, 0.07 mmol) and $K_2CO_3$ (18.5 mg, 0.13 mmol) were added to solution of MeI (4.4 µl, 0.07 mmol) in DMF (1.0 mL) and the resultant mixture was stirred at rt for 16 h. Analysis by LCMS showed residual starting material remaining. A further 0.1 equiv. of MeI in DMF (200 µL) was added, and the reaction mixture left to stir at rt for 1 h, then concentrated under reduced pressure. Purification via normal phase column chromatography (0-90% EtOAc in heptane) gave **5** as an amorphous yellow solid (35 mg, 99% yield).

$^1$H NMR (400 MHz, $CDCl_3$) $\delta$ (ppm) = 7.69 (d, *J* = 8.4 Hz, 1H), 6.92 (dd, *J* = 8.5, 2.2 Hz, 1H), 6.87 – 6.84 (d, *J* = 2.1 Hz, 1H), 5.06 (dd, *J* = 13.4, 5.1 Hz, 1H), 4.31 (d, *J* = 15.9 Hz, 1H), 4.18 (d, *J* = 15.9 Hz, 1H), 4.10 – 4.06 (m, 2H), 4.03 (q, *J* = 7.2 Hz, 2H), 3.76 – 3.73 (m, 2H), 3.70 (t, *J* = 6.4 Hz, 2H), 3.09 (s, 3H), 2.94 – 2.84 (m, 1H), 2.82 – 2.69 (m, 1H), 2.21 (qd, *J* = 13.2, 4.8 Hz, 1H), 2.12 – 2.02 (m, 1H), 1.35 (s, 9H).

**N-(2-methyl-6'-((1-(3-(2-((2-(1-methyl-2,6-dioxopiperidin-3-yl)-1-oxoisoindolin-5-yl)oxy)ethoxy)propanoyl)piperidin-4-yl)oxy)-5'-morpholino-[3,3'-bipyridin]-5-yl)-3-(trifluoromethyl)benzamide, ACBI-0068**

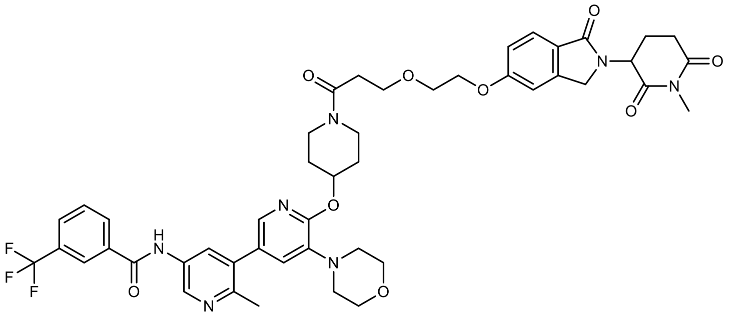

TFA (1.00 mL) was added to a solution of *tert*-butyl 3-(2-((2-(1-methyl-2,6-dioxopiperidin-3-yl)-1-oxoisoindolin-5-yl)oxy)ethoxy)propanoate, **5** (60 mg, 0.13 mmol) in DCM (1.0 mL) at 0°C and the resultant mixture was stirred at rt for 18 h. Analysis by LCMS showed full conversion of the *tert*-butyl ester to the corresponding carboxylic acid. The solvents were removed by aspiration under nitrogen and azeotroped twice with toluene. A mixture of the crude acid, *N*-(2-methyl-5'-morpholino-6'-(piperidin-4-yloxy)-[3,3'-bipyridin]-5-yl)-3-(trifluoromethyl)benzamide hydrochloride, **4** (77.7 mg, 0.13 mmol), HATU (76.6 mg, 0.20 mmol) and DIPEA (109 µl, 0.67 mmol) in DCM (1.0 mL) was stirred at rt for 10 minutes. Analysis by LCMS showed majority conversion to desired product. Purification by normal phase column chromatography (0-10% MeOH in DCM), followed by preparative HPLC purification under acidic conditions (5-95% $CH_3CN$ in 0.1% aq. HCOOH) gave **ACBI-0068** as an amorphous white solid (35 mg, 27% yield).

$^1$H NMR (400 MHz, $CDCl_3$) $\delta$ (ppm) = 8.91 (d, $J$ = 27.8 Hz, 1H), 8.73 (dd, $J$ = 18.6, 2.5 Hz, 1H), 8.16 (d, $J$ = 7.1 Hz, 1H), 8.09 – 8.04 (m, 2H), 7.71 (d, $J$ = 7.8 Hz, 1H), 7.64 – 7.62 (m, 1H), 7.55 (dt, $J$ = 18.8, 7.9 Hz, 2H), 6.96 – 6.94 (m, 1H), 6.90 – 6.78 (m, 2H), 5.34 – 5.28 (m, 1H), 4.95 (dd, $J$ = 13.4, 5.0 Hz, 1H), 4.29 (d, $J$ = 16.0 Hz, 1H), 4.17 (d, $J$ = 16.0 Hz, 1H), 4.09 – 4.04 (m, 2H), 3.86 – 3.70 (m, 9H), 3.68 – 3.61 (m, 1H), 3.57 – 3.50 (s, 2H), 3.46 – 3.38 (m, 1H), 3.05 (s, 3H), 3.03 – 2.98 (m, 4H), 2.88 – 2.80 (m, 1H), 2.71 – 2.51 (m, 3H), 2.41 (s, 3H), 2.25 – 2.09 (m, 1H), 2.08 – 1.88 (m, 3H), 1.81 – 1.74 (m, 1H). HRMS (ESI) for $C_{47}H_{50}F_3N_7O_9$ $[M+H]^+$ calculated 914.3695, obtained 914.3705.

**Synthesis of (*S*)-*N*-(4-methyl-3-(2-morpholino-6-(pyrrolidin-3-yloxy)pyridin-4-yl)phenyl)-2-(trifluoromethyl)isonicotinamide, Intermediate 15**

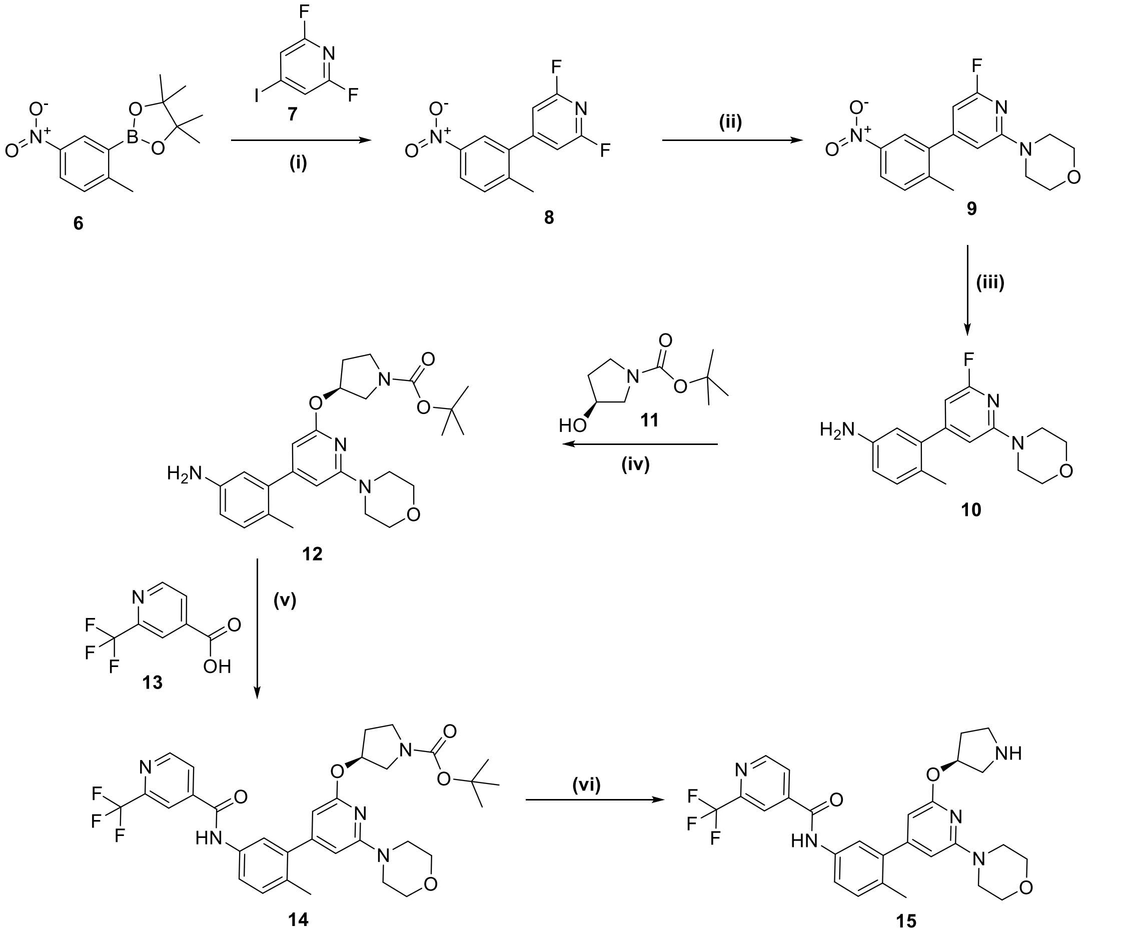

**Synthetic route to intermediate 15.** Reagents and conditions: i) **7**, $Na_2CO_3$, cat. Pd(dppf)$Cl_2$, $H_2O$/DME, 60°C, 6 h; ii) morpholine, DIPEA, EtOH, 70°C, 4 h; iii) 10% Pd/C, EtOH, rt, 5 h; iv) **11**, NaH, dioxane, 110°C, 16 h; v) **13**, HATU, DIPEA, DMF, rt, 2h; vi) 4M HCl in dioxane, dioxane, rt, 2h;

#### 2,6-difluoro-4-(2-methyl-5-nitrophenyl)pyridine, 8

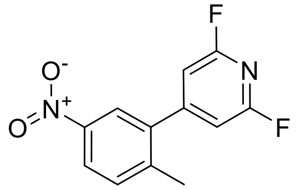

A stirred solution of 4,4,5,5-tetramethyl-2-(2-methyl-5-nitro-phenyl)-[1,3,2]dioxaborolane **6** (5.00 g, 19.0 mmol) in DME (50 mL) was purged with nitrogen for 10 minutes. Sodium carbonate (3.99 g, 38.0 mmol), water (5.0 mL), 2,6-difluoro-4-iodo-pyridine **7** (3.66 g, 15.2 mmol) and Pd(dppf)$Cl_2 \cdot CH_2Cl_2$ (620 mg; 0.760 mmol) were added. The mixture was heated at 60°C for 5 h. The reaction mixture was quenched with water and extracted with EtOAc. The combined organics were dried and concentrated under reduced pressure to give crude **8**, which was taken on to the next step without further purification. MS (ESI) for $C_{12}H_8F_2N_2O_2$ $[M+H]^+$ calculated 251.06, obtained 251.0.

**4-(6-fluoro-4-(2-methyl-5-nitrophenyl)pyridin-2-yl)morpholine, 9**

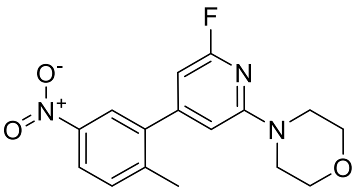

To a stirred solution of 2,6-difluoro-4-(2-methyl-5-nitro-phenyl)-pyridine **8** (6.00 g, 24.0 mmol) in EtOH (100 mL), DIPEA (12.9 mL; 71.9 mmol) was added at 0°C, followed by morpholine (14.62 g; 168 mmol). The resultant mixture was stirred at 70°C for 4 h. The reaction was quenched with cold water, the solid was collected by filtration, rinsed with water, and dried under high vacuum to give crude **9**, which was taken on to the next step without further purification. MS (ESI) for $C_{16}H_{16}FN_3O_3$ $[M+H]^+$ calculated 318.12, obtained 318.1.

**3-(2-fluoro-6-morpholinopyridin-4-yl)-4-methylaniline, 10**

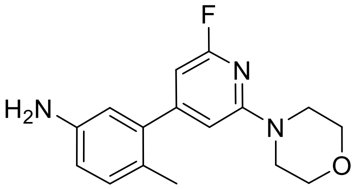

To a stirred solution of 4-(6-Fluoro-4-(2-methyl-5-nitro-phenyl)-pyridin-2-yl)-morpholine **9** (6.00 g, 18.9 mmol) in EtOH (400 mL), 10% Pd/C (4.80 g, 4.53 mmol) was added portion wise, and the mixture was hydrogenated in a steel reactor at 50-60 PSI for 4 h. The reaction mixture was filtered through a Celite pad and rinsed with MeOH. The filtrate was concentrated under reduced pressure. Purification via column chromatography (neutral alumina, 0-50% EtOAc in petroleum ether) gave **10** as an amorphous white solid (4.50 g, 83% yield).

$^1$H NMR (500 MHz, $CDCl_3$) δ 7.04 (d, *J* = 8.1 Hz, 1H), 6.65 (dd, *J* = 8.1, 2.6 Hz, 1H), 6.54 (d, *J* = 2.6 Hz, 1H), 6.33 (d, *J* = 1.4 Hz, 1H), 6.18 (d, *J* = 1.4 Hz, 1H), 3.81 (dd, *J* = 5.8, 4.1 Hz, 4H), 3.62 (s, 2H), 3.51 (dd, *J* = 5.8, 4.1 Hz, 4H), 2.15 (s, 3H). MS (ESI) for $C_{16}H_{18}FN_3O$ $[M+H]^+$ calculated 288.14, obtained 288.1.

***tert*-butyl (*S*)-3-((4-(5-amino-2-methylphenyl)-6-morpholinopyridin-2-yl)oxy)pyrrolidine-1-carboxylate, 12**

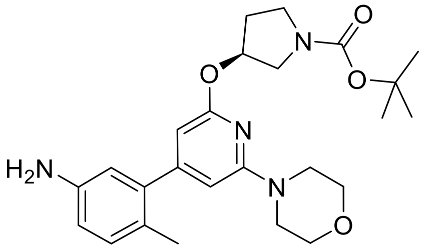

NaH (1.11 g, 27.9 mmol) was added portion wise to a solution of *tert*-butyl (S)-3-hydroxypyrrolidine-1-carboxylate **11** (4.17 g, 22.3 mmol) in dioxane (32 mL) and stirred at rt for 30 min. A solution of 3-(2-fluoro-6-morpholinopyridin-4-yl)-4-methylaniline, **10** (3.20 g, 11.1 mmol) was then added and the resultant mixture was stirred at 110°C for 16 h. The reaction mixture was quenched with water and extracted with EtOAc. The combined organics were dried and concentrated under reduced pressure to give crude **12**, which was taken on to the next step without further purification. MS (ESI) for $C_{25}H_{34}N_4O_4$ $[M+H]^+$ calculated 455.26, obtained 455.6.

***tert*-butyl (*S*)-3-((4-(2-methyl-5-(2-(trifluoromethyl)isonicotinamido)phenyl)-6-morpholinopyridin-2-yl)oxy)pyrrolidine-1-carboxylate, 14**

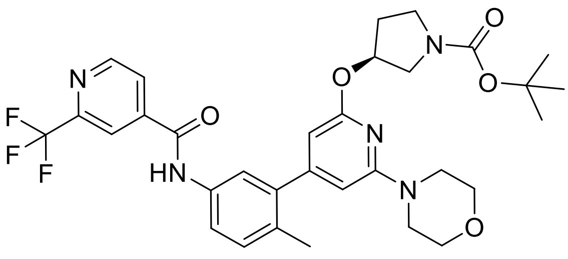

HATU (1.25 g, 3.30 mmol), DIPEA (1.15 mL, 6.60 mmol) and *tert*-butyl (*S*)-3-((4-(5-amino-2-methylphenyl)-6-morpholinopyridin-2-yl)oxy)pyrrolidine-1-carboxylate **12** (1.00 g, 2.20 mmol) were added to a stirred solution of 2-(trifluoromethyl)isonicotinic acid **13** (420 mg; 2.20 mmol) in DMF (10 mL) and the resultant mixture was stirred at rt for 2 h. The reaction mixture was quenched with cold water and the precipitated solid was filtered, rinsed with water, then dried at high vacuum to obtain crude **14**, which was taken on to the next step without further purification. MS (ESI) for $C_{32}H_{36}F_3N_5O_5$ $[M+H]^+$ calculated 628.27, obtained 628.8.

**(*S*)-*N*-(4-methyl-3-(2-morpholino-6-(pyrrolidin-3-yloxy)pyridin-4-yl)phenyl)-2-(trifluoromethyl) isonicotinamide, 15**

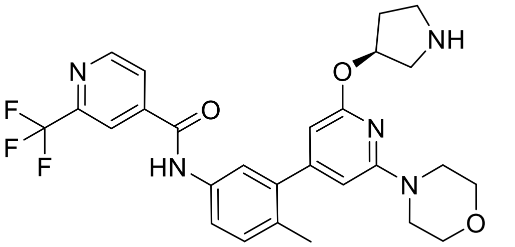

4 M HCl in dioxane (10.0 mL, 40.0 mmol) was added to a stirred solution of *tert*-butyl (*S*)-3-((4-(2-methyl-5-(2-(trifluoromethyl)isonicotinamido)phenyl)-6-morpholinopyridin-2-yl)oxy)pyrrolidine-1-carboxylate **14** (1.50 g, 2.39 mmol) in DCM (15.0 mL) at 0°C and the resultant mixture was stirred at rt for 2 h. The solvent was evaporated under reduced pressure. Purification by preparative HPLC under basic conditions (5-95% $CH_3CN$ in 0.1% aq. $NH_3$) gave **15** as an amorphous brown solid (800 mg, 63% yield).

$^1H$ NMR (500 MHz, $CDCl_3$), (n.b. doubling seen for many peaks due to presence of rotamers, *minor rotamer): δ 8.89 – 8.83 (m, 1H), 8.59 (br s, 1H), 8.16 (s, 0.8H), 8.13 (s, 0.2H)*, 7.97 (d, *J* = 5.0 Hz, 0.8H), 7.93 (d, *J* = 5.0 Hz, 0.2H)*, 7.69 – 7.62 (m, 1H), 7.41 – 7.35 (m, 1H), 7.24 (d, *J* = 8.4 Hz, 1H), 6.08 (s, 0.85H), 6.03 (app s, 0.3H)*, 5.99 (s, 0.85H), 5.46 (s, 0.85H), 5.41 (s, 0.15H)*, 3.80 (t, *J* = 4.9 Hz, 4H), 3.46 (t, *J* = 5.0 Hz, 4H), 3.30 – 3.17 (m, 3H), 3.07 (app s, 1H), 2.24 (s, 3H), 2.21 – 2.10 (m, 1H), 2.13 – 2.04 (m, 1H). MS (ESI) for $C_{27}H_{28}F_3N_5O_3$ $[M+H]^+$ calculated 528.21, obtained 528.4

#### Synthesis of *N*-(3-(2-(((3*S*)-1-(3-(2-((2-(2,6-dioxopiperidin-3-yl)-1,3-dioxoisoindolin-5-yl)oxy)ethoxy)propanoyl)pyrrolidin-3-yl)oxy)-6-morpholinopyridin-4-yl)-4-methylphenyl)-2-(trifluoromethyl)isonicotinamide, ACBI-5767

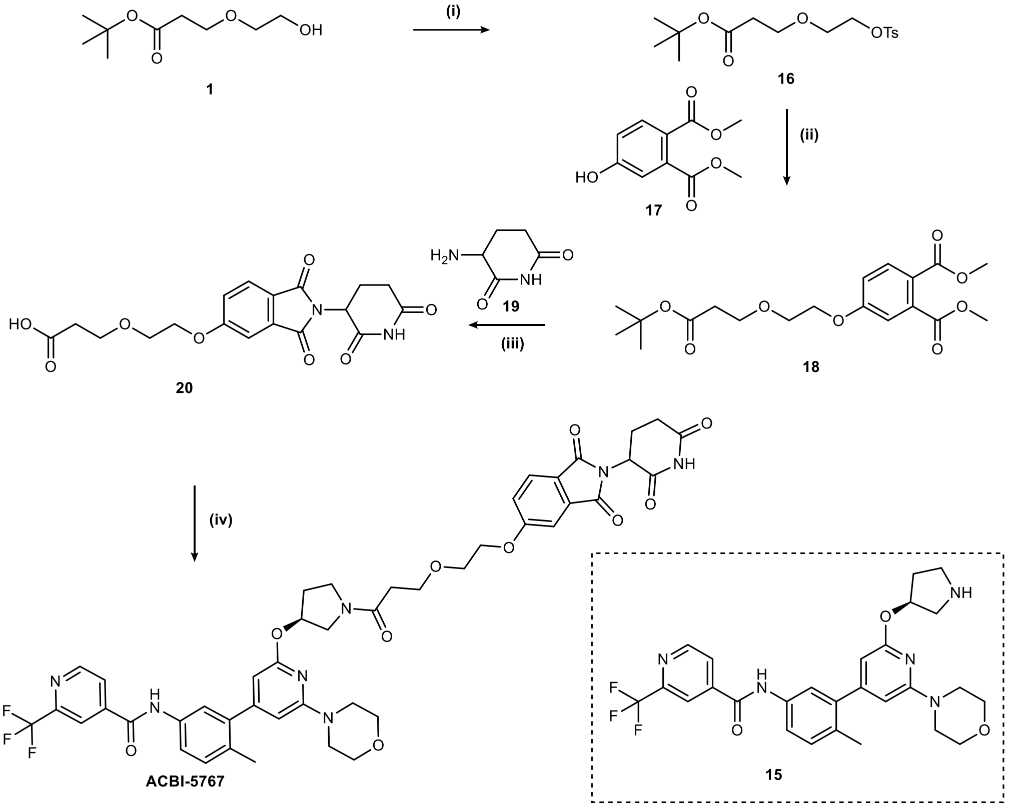


**Synthetic route to compound ACBI-5767.** Reagents and conditions: i) TsCl, DMAP, $Et_3N$, $CH_2Cl_2$, rt, 18 h; ii) **17**, $K_2CO_3$, DMF, 60°C, 18 h; iii) **19**, 4M aq NaOH, Pyridine, MeOH, 70°C, 5 h; iv) **15**, HATU, DIPEA, MeCN, rt, 18 h.

##### *tert*-butyl 3-(2-(tosyloxy)ethoxy)propanoate, 16

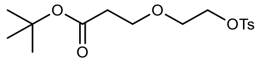

Tosyl Chloride (782 mg, 4.15 mmol) in DCM (3.0 mL) was added to a mixture of *tert*-butyl 3-(2-hydroxyethoxy)propanoate **1** (608 mg, 3.19 mmol), $Et_3N$ (580 µL, 4.15 mmol) and DMAP (117 mg, 0.95 mmol) in DCM (4.0 mL) at 0°C and the resultant mixture was stirred at rt for 18 h. Sat. aq. $NaHCO_3$ (5 mL) was added and the mixture was extracted with DCM. The combined organics are washed with brine, then dried and concentrated under reduced pressure. Purification via column chromatography (0-50% EtOAc in hexane) gave **16** as a white amorphous solid (907 mg, 66% yield).
$^1$H NMR (500 MHz, $CDCl_3$) $\delta$ (ppm) = 7.73 (d, *J* = 8.3 Hz, 2H), 7.27 (d, *J* = 8.0 Hz, 2H), 4.12 – 4.03 (m, 2H), 3.63 – 3.51 (m, 4H), 2.38 (s, 3H), 2.34 (t, *J* = 6.4 Hz, 2H), 1.37 (s, 9H).

##### Dimethyl 4-(2-(3-(tert-butoxy)-3-oxopropoxy)ethoxy)phthalate, 18

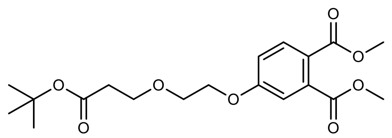

Dimethyl 4-hydroxyphthalate **17** (400 mg, 1.9 mmol) was added to *tert*-butyl 3-(2-(tosyloxy)ethoxy)propanoate **16** (787 mg, 2.28 mmol) in DMF (4.0 mL) and the resultant mixture was stirred at 60°C for 18 h. The reaction mixture was diluted with water, DCM was added and the mixture passed through a phase separator. The organics were concentrated under reduced pressure.

Purification via column chromatography (0-100% EtOAc in heptane) gave **18** as an amorphous yellow solid (209 mg, 27% yield).
$^1$H NMR (500 MHz, $CDCl_3$) *δ* (ppm) = 7.81 (d, *J* = 8.7 Hz, 1H), 7.12 (d, *J* = 2.6 Hz, 1H), 7.03 (dd, *J* = 8.7, 2.6 Hz, 1H), 4.19 (dd, *J* = 5.6, 3.9 Hz, 2H), 3.93 (s, 3H), 3.89 (s, 3H), 3.87 – 3.83 (m, 2H), 3.81 (t, *J* = 6.4 Hz, 2H), 2.54 (t, *J* = 6.4 Hz, 2H), 1.47 (s, 9H).

**3-(2-((2-(2,6-dioxopiperidin-3-yl)-1,3-dioxoisoindolin-5-yl)oxy)ethoxy)propanoic acid, 20**

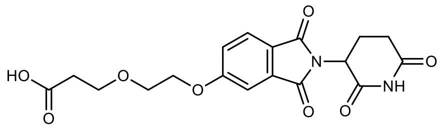


A mixture of 3-aminopiperidine-2,6-dione hydrochloride **19** (81 mg, 0.49 mmol), dimethyl 4-(2-(3-(tert-butoxy)-3-oxopropoxy)ethoxy)phthalate **18** (209 mg, 0.55 mmol), and NaOH (4M aq., 400 µL, 1.58 mmol) in pyridine (1.5 mL, 18.6 mmol) was heated at 70°C for 5 h. The reaction mixture was diluted with water and extracted with EtOAc. The combined organics were dried and concentrated under reduced pressure. Purification via column chromatography (0-100% EtOAc in heptane) gave **20** as a yellow oil (74 mg, 33% yield, 80% purity).
$^1$H NMR (500 MHz, MeOD) δ 6.34 (d, *J* = 8.3 Hz, 1H), 5.97 (d, *J* = 2.3 Hz, 1H), 5.88 (dd, *J* = 8.3, 2.3 Hz, 1H), 3.64 (dd, *J* = 12.4, 5.5 Hz, 1H), 2.91 – 2.80 (m, 2H), 2.46 – 2.39 (m, 2H), 2.36 (t, *J* = 6.2 Hz, 2H), 1.41 (ddd, *J* = 17.9, 14.1, 5.2 Hz, 1H), 1.34 – 1.27 (m, 2H), 1.12 (t, *J* = 6.2 Hz, 2H), 0.71 – 0.62 (m, 1H). MS (ESI) for $C_{18}H_{18}N_2O_8$ $[M+H]^+$ calculated 391.11, obtained 390.9.

***N*-(3-(2-(((3S)-1-(3-(2-((2-(2,6-dioxopiperidin-3-yl)-1,3-dioxoisoindolin-5-yl)oxy)ethoxy)propanoyl)pyrrolidin-3-yl)oxy)-6-morpholinopyridin-4-yl)-4-methylphenyl)-2-(trifluoromethyl)isonicotinamide, ACBI-5767**

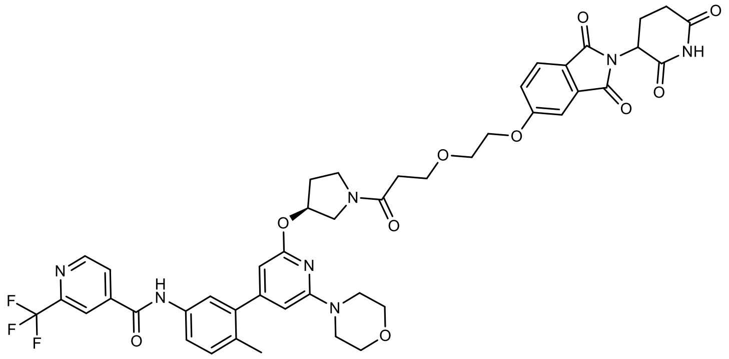


DIPEA (52 µL, 0.30 mmol) was added to a solution of 3-(2-((2-(2,6-dioxopiperidin-3-yl)-1,3-dioxoisoindolin-5-yl)oxy)ethoxy)propanoic acid, **20** (39 mg, 0.10 mmol), (*S*)-*N*-(4-methyl-3-(2-morpholino-6-(pyrrolidin-3-yloxy)pyridin-4-yl)phenyl)-2-(trifluoromethyl) isonicotinamide, **15** (53 mg, 0.10 mmol), and HATU (57 mg, 0.15 mmol) in MeCN (2.0 mL). The resultant mixture was stirred at rt for 18 h, then concentrated under reduced pressure. Purification by preparative HPLC under acidic conditions (5-95% $CH_3CN$ in 0.1% aq. HCOOH) gave **ACBI-5767** as an amorphous white solid (18 mg, 19% yield).
$^1$H NMR (500 MHz, $CDCl_3$) *δ* (ppm) = 8.82 (d, *J* = 5.0 Hz, 1H), 8.34 – 8.28 (m, 1H), 8.05 (s, 1H), 8.05 – 7.98 (m, 1H), 7.87 (d, *J* = 5.0 Hz, 1H), 7.58 (td, *J* = 7.8, 2.3 Hz, 1H), 7.48 – 7.42 (m, 1H), 7.38 (d, *J* = 18.6 Hz, 1H), 7.22 – 7.20 (m, 1 Hz, 1H), 7.18 – 7.15 (m, 1H), 7.10 – 7.06 (m, 1H), 6.02 (d, *J* = 14.6 Hz, 1H), 5.98 – 5.87 (m, 1H), 5.57 – 5.41 (m, 1H), 4.81 – 4.86 (m, 1H), 4.11 (dt, *J* = 9.5, 4.8 Hz, 2H), 3.84 – 3.79 (m, 2H), 3.79 – 3.69 (m, 6H), 3.68 – 3.52 (m, 4H), 3.42 – 3.37 (m, 4H), 2.84 – 2.78 (m, 1H), 2.76 – 2.62 (m, 2H), 2.60 – 2.44 (m, 2H), 2.27 – 2.21 (m, 1H), 2.18 – 2.15 (m, 4H), 2.06 – 2.02 (m, 1H). HRMS (ESI) for $C_{45}H_{45}F_3N_7O_{10}$ $[M+H]^+$ calculated 900.3175, 900.3196.

### NMR and HRMS SPECTRA OF ACBI-8451, ACBI-0068, ACBI-5767

#### ACBI-8451

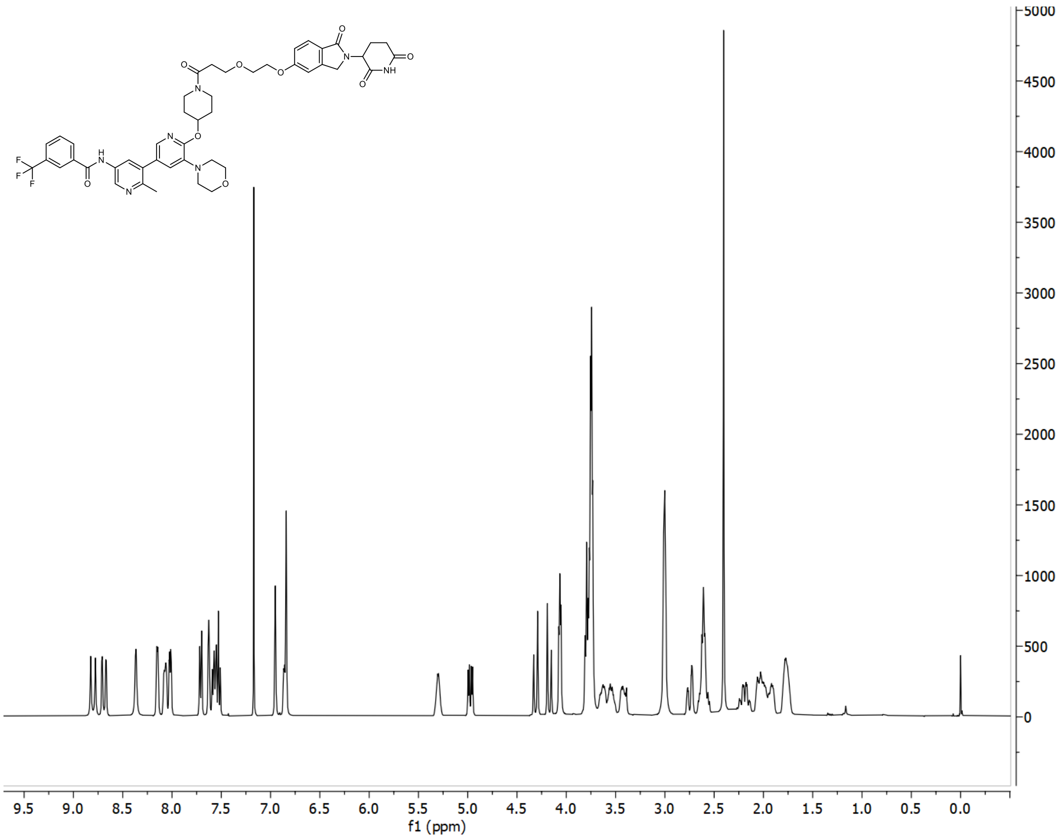


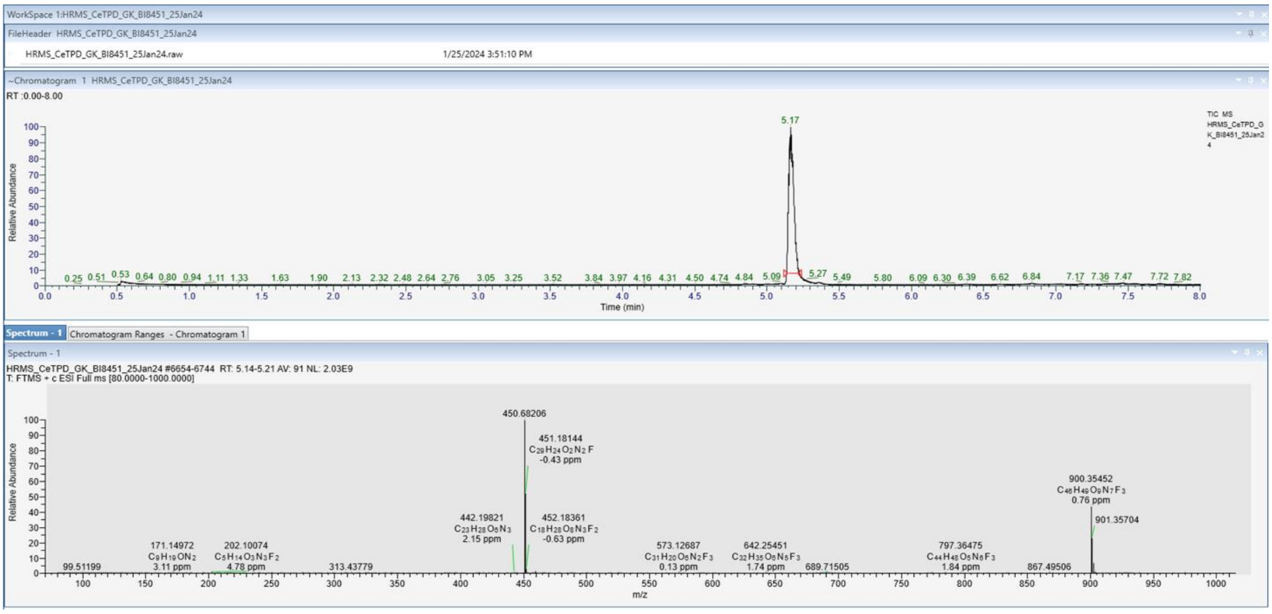


#### ACBI-0068

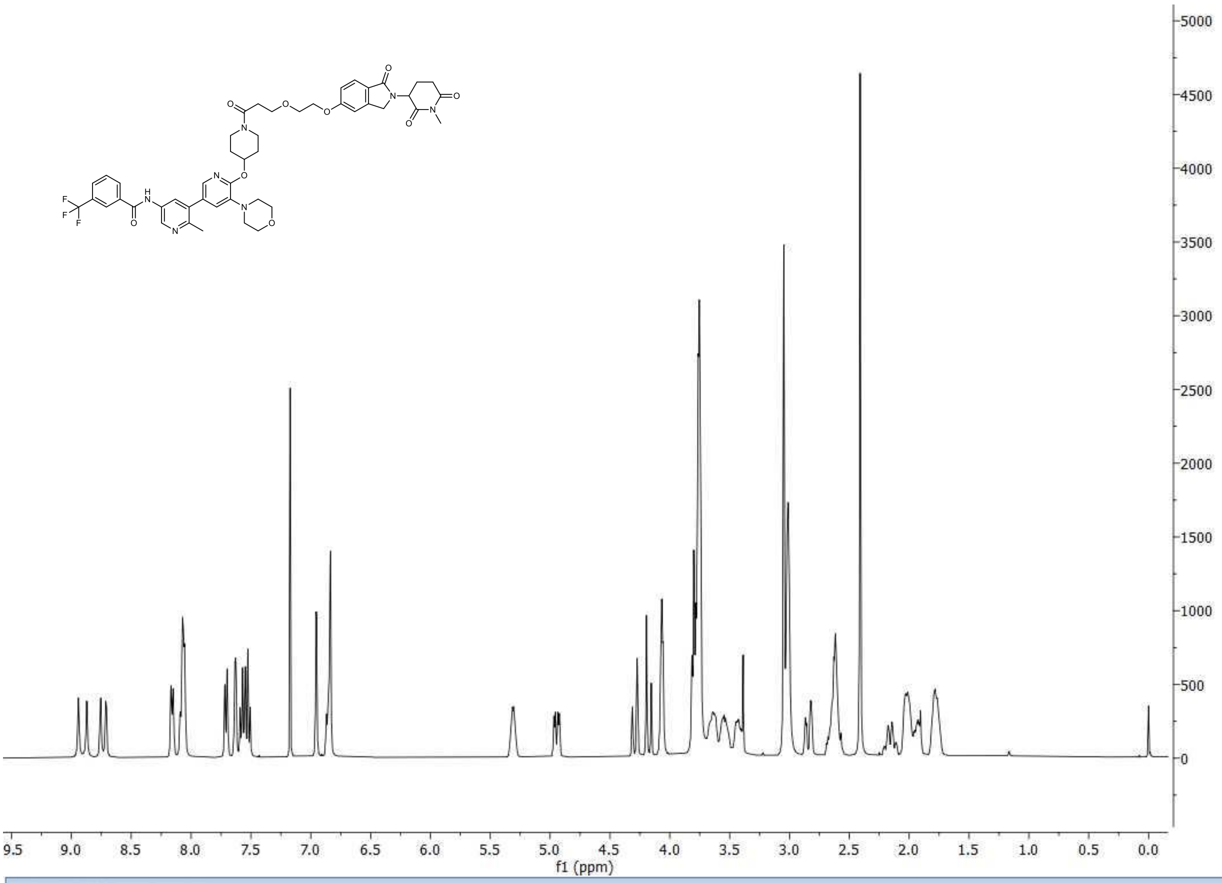

f1 (ppm)


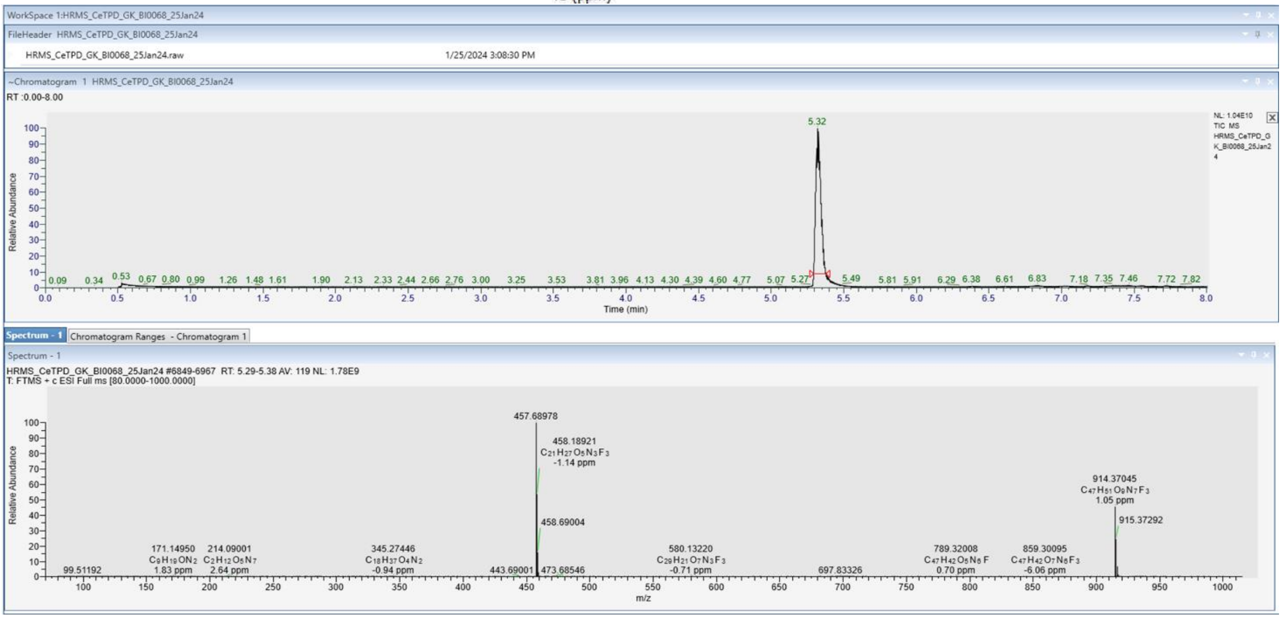

WorkSpace 1:HRMS_CeTPD_GK_BI0068_25Jan24
FileHeader HRMS_CeTPD_GK_BI0068_25Jan24
HRMS_CeTPD_GK_BI0068_25Jan24.raw
1/25/2024 3:08:30 PM
~Chromatogram 1 HRMS_CeTPD_GK_BI0068_25Jan24
RT :0.00-8.00
NL: 1.04E10
TIC MS
HRMS_CeTPD_GK_BI0068_25Jan24
Relative Abundance
Time (min)
5.32
Spectrum - 1
Chromatogram Ranges - Chromatogram 1
HRMS_CeTPD_GK_BI0068_25Jan24 #6849-6967 RT: 5.29-5.38 AV: 119 NL: 1.78E9
T: FTMS + c ESI Full ms [80.0000-1000.0000]
457.68978
458.18921
C21 H27 O5 N3 F3
-1.14 ppm
458.69004
914.37045
C47 H51 O9 N7 F3
1.05 ppm
915.37292
m/z

#### ACBI-5767

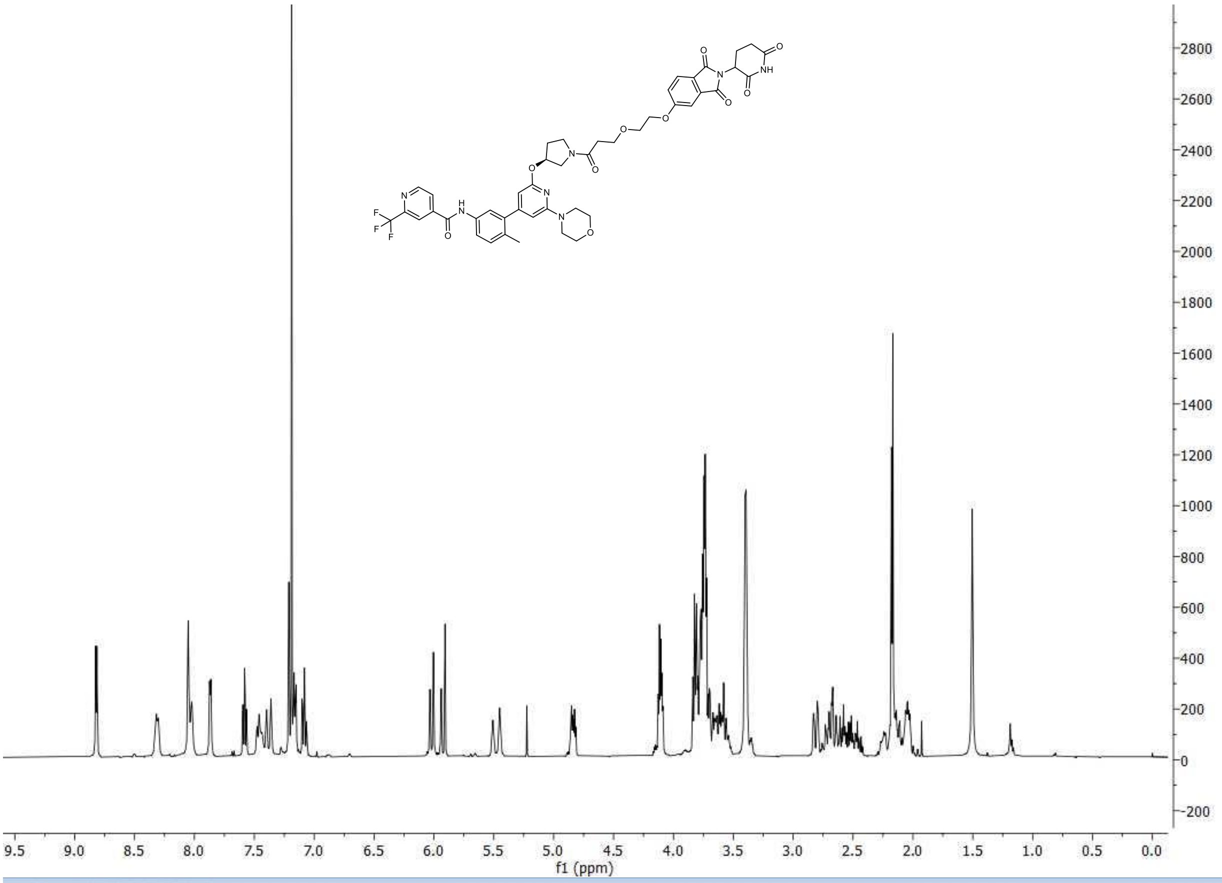

f1 (ppm)


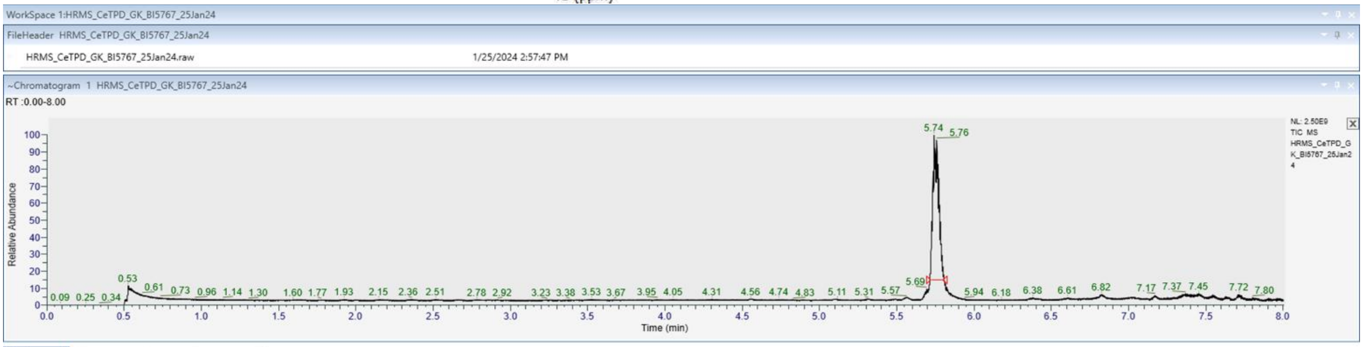

WorkSpace 1:HRMS_CeTPD_GK_BI5767_25Jan24
FileHeader HRMS_CeTPD_GK_BI5767_25Jan24
HRMS_CeTPD_GK_BI5767_25Jan24.raw
1/25/2024 2:57:47 PM
~Chromatogram 1 HRMS_CeTPD_GK_BI5767_25Jan24
RT :0.00-8.00
Relative Abundance
Time (min)
5.74
5.76


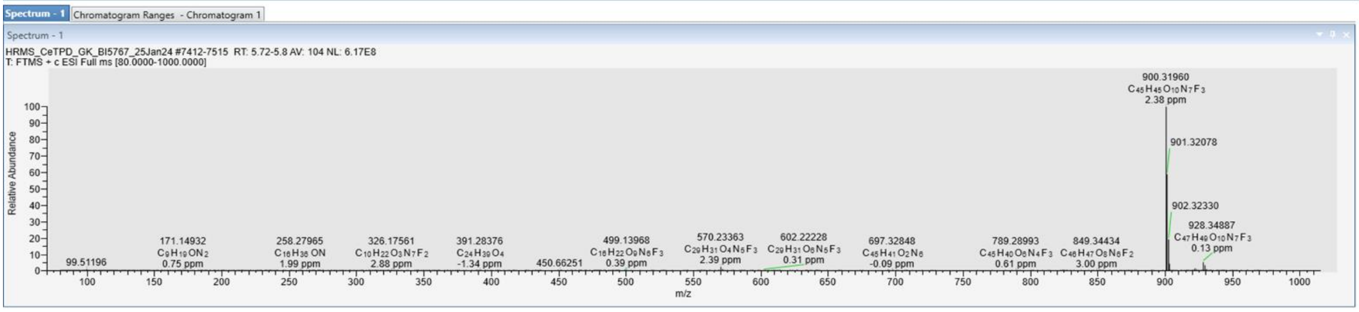

Spectrum - 1
Chromatogram Ranges - Chromatogram 1
HRMS_CeTPD_GK_BI5767_25Jan24 #7412-7515 RT: 5.72-5.8 AV: 104 NL: 6.17E8
T: FTMS + c ESI Full ms [80.0000-1000.0000]
900.31960
C45 H45 O10 N7 F3
2.38 ppm
901.32078
902.32330
928.34887
C47 H49 O10 N7 F3
0.13 ppm
Relative Abundance
m/z
